# Antiviral Functions of ARGONAUTE Proteins During *Turnip Crinkle Virus* Infection Revealed by Image-based Trait Analysis in *Arabidopsis*

**DOI:** 10.1101/487322

**Authors:** Xingguo Zheng, Noah Fahlgren, Arash Abbasi, Jeffrey C. Berry, James C. Carrington

## Abstract

RNA-based silencing functions as an important antiviral immunity mechanism in plants. Plant viruses evolved to encode viral suppressors of RNA silencing (VSRs) that interfere with the function of key components in the silencing pathway. As effectors in the RNA silencing pathway, ARGONAUTE (AGO) proteins are targeted of by some VSRs, such as that encoded by *Turnip crinkle virus* (TCV). A VSR-deficient TCV mutant was used to identify AGO proteins with antiviral activities during infection. A quantitative phenotyping protocol using an image-based color trait analysis pipeline on the PlantCV platform, with temporal red, green and blue (RGB) imaging and a computational segmentation algorithm, was used to measure plant disease after TCV inoculation. This process captured and analyzed growth and leaf color of *Arabidopsis* plants in response to virus infection over time. By combining this quantitative phenotypic data with molecular assays to detect local and systemic virus accumulation, AGO2, AGO3, and AGO7 were shown to play antiviral roles during TCV infection. In leaves, AGO2 and AGO7 functioned as prominent non-additive, anti-TCV effectors, while AGO3 played a minor role. Other AGOs were required to protect inflorescence tissues against TCV. Overall, these results indicate that distinct AGO proteins have specialized, modular roles in antiviral defense across different tissues, and demonstrate the effectiveness of image-based phenotyping to quantify disease progression.

**Author Summary:** Plant viruses caused substantial losses in crop production and quality worldwide. Precisely measuring plant health is critical for better understanding the mechanisms underlying plant virus and host interactions. Advances in high-resolution imaging technologies and deep-learning tools have made acquiring and analyzing “big data” of disease traits possible. In this study, we have developed a high-throughput, image-based trait phenotyping pipeline to quantify disease severity in *Arabidopsis thaliana* infected by *Turnip Crinkle Virus* (TCV). Our aim is to understand how the antiviral RNA silencing machinery is tuned to protect the host from invading virus infection. We focused on ARGONAUTE proteins, which are the effectors in the RNA silencing pathway. A mutant line of TCV with a dysfunctional silencing suppressor (P38) was used to investigate which *ago* mutation could compensate for the dysfunctional silencing suppressor and facilitate the development of disease symptoms. We demonstrated that specific AGO proteins contribute to protecting leaves from TCV infection in a non-additive manner. Our results also implied that distinct AGOs are required to function collectively to silence TCV in inflorescence tissues. More evidence is still needed to further understand how these antiviral AGOs interact with suppressor proteins molecularly during TCV infection.

## Introduction

Plants can protect themselves against invasive virus infection through RNA silencing by targeting viral RNA for degradation [1]. This host silencing machinery is triggered by viral double-stranded RNAs (dsRNAs), which are cleaved by Dicer-like (DCL) nucleases associated with dsRNA binding proteins (DRBs) into 21–24 nucleotide RNA duplexes called viral small interfering RNAs (vsiRNAs). The vsiRNAs are then methylated and stabilized by HUA enhancer 1 (HEN1). One strand of these stabilized vsiRNAs is recruited into the RNA-induced silencing complex (RISC) containing ARGONAUTE (AGO) proteins, and then serves as the sequence-specific guide for specific AGOs to slice cognate viral RNAs [2, 3].

Most plant viruses have evolved to encode viral suppressors of RNA silencing (VSRs) that use varied mechanisms to target components in the silencing pathway [4]. One such mechanism is interference of AGOs that mediate antiviral silencing. Accumulating evidence indicates that various VSRs use diverse modes of action on AGO proteins, such as promoting AGO degradation [5–8], inhibiting the slicing activities of AGOs [9], or interfering with factors upstream of AGO activity such as RNA-dependent RNA Polymerase (RDR)-dependent silencing [10], obstructing siRNA-loaded RISC activity [11, 12], or indirectly repressing AGO protein level [13]. Since functional VSRs can mask host antiviral silencing effects, VSR-defective mutant viruses that can only successfully infect immunocompromised plants have been constructed to identify antiviral roles of key components in the silencing pathway during virus infection [14, 15].

The *Arabidopsis thaliana* (*Arabidopsis*) genome encodes ten AGO proteins, several of which have demonstrated antiviral roles [16]. AGO1 [14, 17–19] and AGO2 [15, 20–22] have been identified as prominent antiviral AGOs against several RNA viruses, while other AGOs, including AGO5, AGO7, and AGO10, have limited antiviral roles in some cases [14, 15]. For DNA viruses, AGO4 is the main effector protein in the antiviral silencing machinery [23, 24]. The full complement of AGOs with roles in antiviral defense and how distinct antiviral AGOs are coordinated to silence different viruses in different tissues remain to be fully determined.

*Turnip crinkle* virus (TCV) is a positive single-strand RNA virus belonging to the *Carmovirus* genus of the *Tombusviridae* family. The TCV genome encodes five proteins, including two replicase proteins (P28 and P88), two movement proteins (P8 and P9) and the coat protein (P38). The coat protein (CP) is multifunctional, as it has roles in virus movement [25, 26], serves as a virulence factor [27], and functions as a VSR to suppress antiviral silencing [28, 29]. Other virus groups in the *Tombusviridae* encode separate VSR proteins, such as P19 from *Tomato bushy stunt virus* (TBSV) [19]. TCV systemically infects the susceptible *Arabidopsis* ecotype *Columbia-0 (Col-0),* and causes disease symptoms that include severe chlorosis in leaves, stunted bolts and reduced biomass. A previous study reported that replacement of a single amino acid residue in the TCV CP P38 (R130T) disrupts its VSR function without affecting other functions [26]. The VSR-deficient TCV is unable to suppress the host antiviral silencing machinery, leading to a lack of disease symptoms in wild-type plants post-inoculation.

In this study, the roles of *Arabidopsis* AGO proteins in anti-TCV silencing were analyzed using genetic and image-based quantitative phenotyping approaches. Most previous pathological studies have relied on qualitative and subjective visual scoring systems to identify and assess disease phenotypes in plants [30]. A machine learning method [31] and other analysis tools in the open-source PlantCV platform [32, 33] were used to develop to detect subtle, reproducible differences in disease symptoms over time.

## Result

### Suppressor-deficient TCV is not able to elicit disease symptoms in wild-type host plants

*Turnip Crinkle Virus* (TCV) and other carmoviruses use their coat proteins (CPs) as VSRs to suppress host antiviral silencing [34, 35]. By replacing a single amino acid in TCV CP (P38) with its counterpart residue in TBSV CP (R130T) (Fig 1A), the VSR function of TCV CP (P38) is abolished [26]. Introducing this mutation (R130T) in the coat protein did not affect its ability to assemble functional virion particles (S1 Fig). We confirmed that this mutant virus (TCV CPB) lost its capacity to suppress host antiviral silencing by quantifying and comparing the effects of TCV CPB in wild-type *(Col-0)* and *dcl2-1 dcl3-2 dcl4-2 (dcl2 dcl3 dcl4)* triple mutant plants. In the *dcl* triple mutant, three *DCL* genes with roles in antiviral defense are dysfunctional, so it serves as a hyper-susceptible control genotype. Parental TCV infection caused severe chlorosis in *Arabidopsis* leaves in both wild-type control *(Col-0)* and the *dcl* triple mutant plants, while TCV CPB caused similar chlorosis symptoms only in the *dcl* triple mutant (Fig 1B).

**Fig 1.**
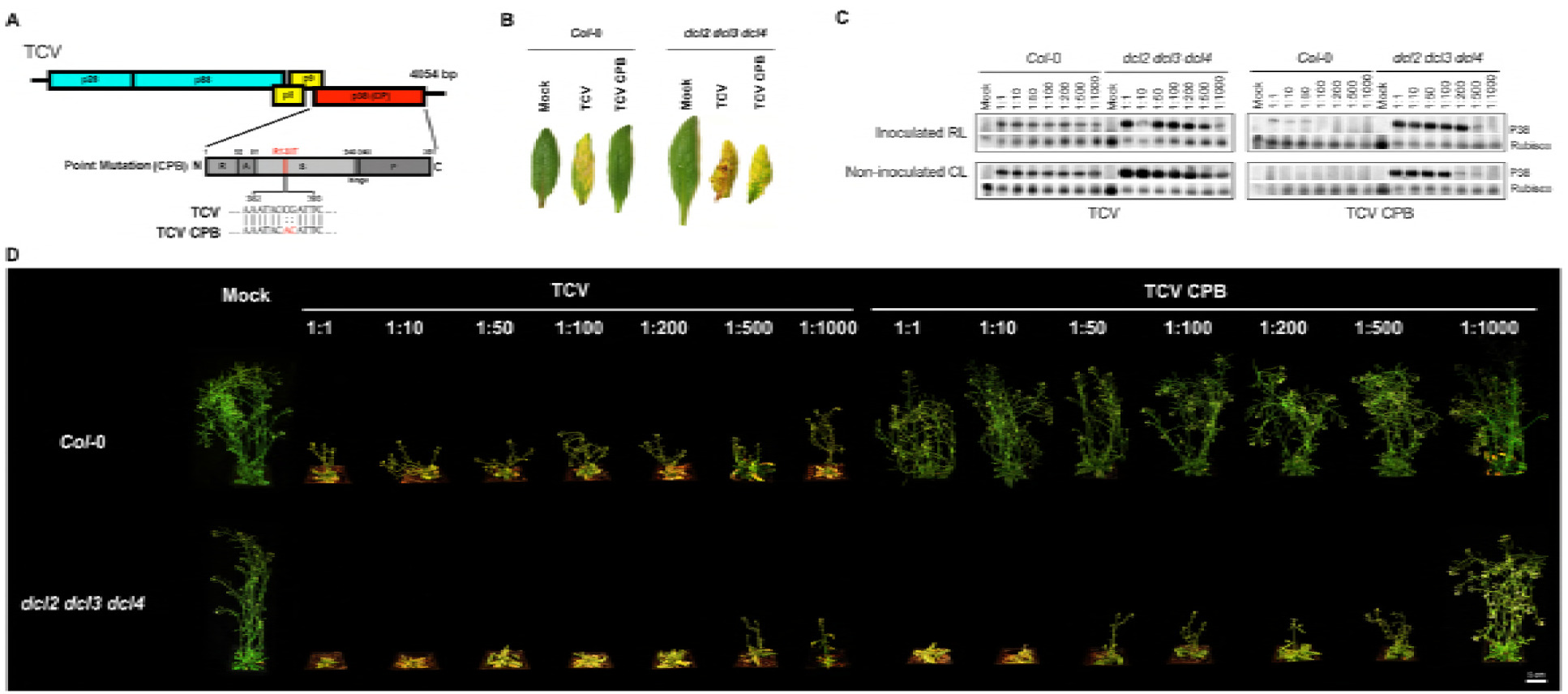
TCV infection-caused disease symptoms in *A. thaliana.* (A) Schematic representation of the TCV genome showing the CPB point mutation on the coat protein (P38). Top diagram shows the genomic RNA of TCV. The bottom diagram shows the coat protein (P38) region of single-amino-acid mutant TCV CPB. The single amino acid change in TCV CPB mutant is marked beneath the bottom diagram. The bottom diagram represents the full-length CP, with the sizes and the relative positions of the five structural domains R: RNA-binding domain; A: arm domain; S: surface domain; hinge; P: protruding domain. (B) TCV infection-caused chlorosis in non-inoculated cauline leaves. Left panel: wild type control *(Col-0);* right panel: hyper-susceptible control *(dcl2 dcl3 dcl4).* Photographs were taken at 14 days post inoculation (dpi). (C) Local and systemic accumulation of coat protein (P38) caused by TCV (left panel) and TCV CPB (right panel) infection was assayed by immunoblotting. P38 was detected using anti-P38 antibody. Rubisco protein was detected by anti-Rubisco antibody as a control. The virion inoculum was in different dilutions. RL: rosette leaf; CL: cauline leaf. RL samples were collected at 7 dpi; CL samples were collected at 14 dpi. (D) TCV infection-caused stunt bolt phenotype in Arabidopsis. Upper panel: wild type control *(Col-0);* bottom panel: hyper-susceptible control *(dcl2 dcl3 dcl4).* The virion inoculum was in different dilutions. Photographs were taken at 14 dpi.

*Turnip crinkle virus* was from samples of inoculated rosette leaves and non-inoculated cauline leaves using immunoblot assays with CP antisera (anti-P38). High levels of CP were detected in rosette leaves at 7 days post inoculation (dpi) and in cauline leaves at 14 dpi of both *Col-0* and *dcl2 dcl3 dcl4* plants infected with parental TCV, even at 1:1000 dilution (Fig 1C). In contrast, low levels of CP were detected in rosette leaves of wild-type control plants inoculated with TCV CPB at lower dilutions (1:1 to 1:50) but not at higher dilutions (1:100 to 1:1000; Fig 1C). In non-inoculated cauline leaves of *Col-0* plants, no observable CPB-P38 signal was detected at any dilution (Fig 1C). However, TCV CPB inoculation of *dcl2 dcl3 dcl4* mutant led to high levels of CP protein accumulation in the rosette and cauline leaves at each dilution ranging from 1:1 to 1:500 (Fig 1C). Notably, in *dcl2 dcl3 dcl4* mutants, no observable local or systemic CPB CP protein was detected when the TCV CPB inoculum was highly diluted (1:1000) (Fig 1C).

Parental TCV infection at any dilution had negative effects on the growth of both *Col-0* and *dcl2 dcl3 dcl4* plants (Fig 1D). Morphologically, plants infected with TCV were shorter compared to non-infected plants (Fig 1D). In contrast, the TCV CPB mutant affected growth of only *dcl2 dcl3 dcl4* plants (Fig 1D). These results confirmed that the CPB (R130T) mutation in TCV CP attenuates VSR functions, and suggested that TCV CPB could be used in a genetic analysis to identify components of the silencing machinery necessary for antiviral defense.

### Image-based analysis of disease symptoms

Before initiating a systematic screen of *Arabidopsis ago* mutants using TCV and TCV CPB, a non-destructive, computer vision-based system using the PlantCV platform [33] was developed for high-resolution, quantitative assessment of disease symptom phenotypes over time (Fig 2A). The system was designed to identify individual plant leaves, measure their areas, lengths, and pixel color characteristics, and distinguish subtle differences in responses of plants with different genotypes over time. Top-down RGB images of individual plants were captured every other day, from one day pre-inoculation to seventeen days post-inoculation. The images were analyzed using a machine learning approach to segment plant from background and to classify plant pixels as “healthy” pixels (green color) or “unhealthy” chlorotic/necrotic pixels (any non-green color; Fig 2B). The ratio of chlorotic/necrotic to total plant pixels was used to calculate the extent and severity of symptomatic rosette leaf tissue.

**Fig 2.**
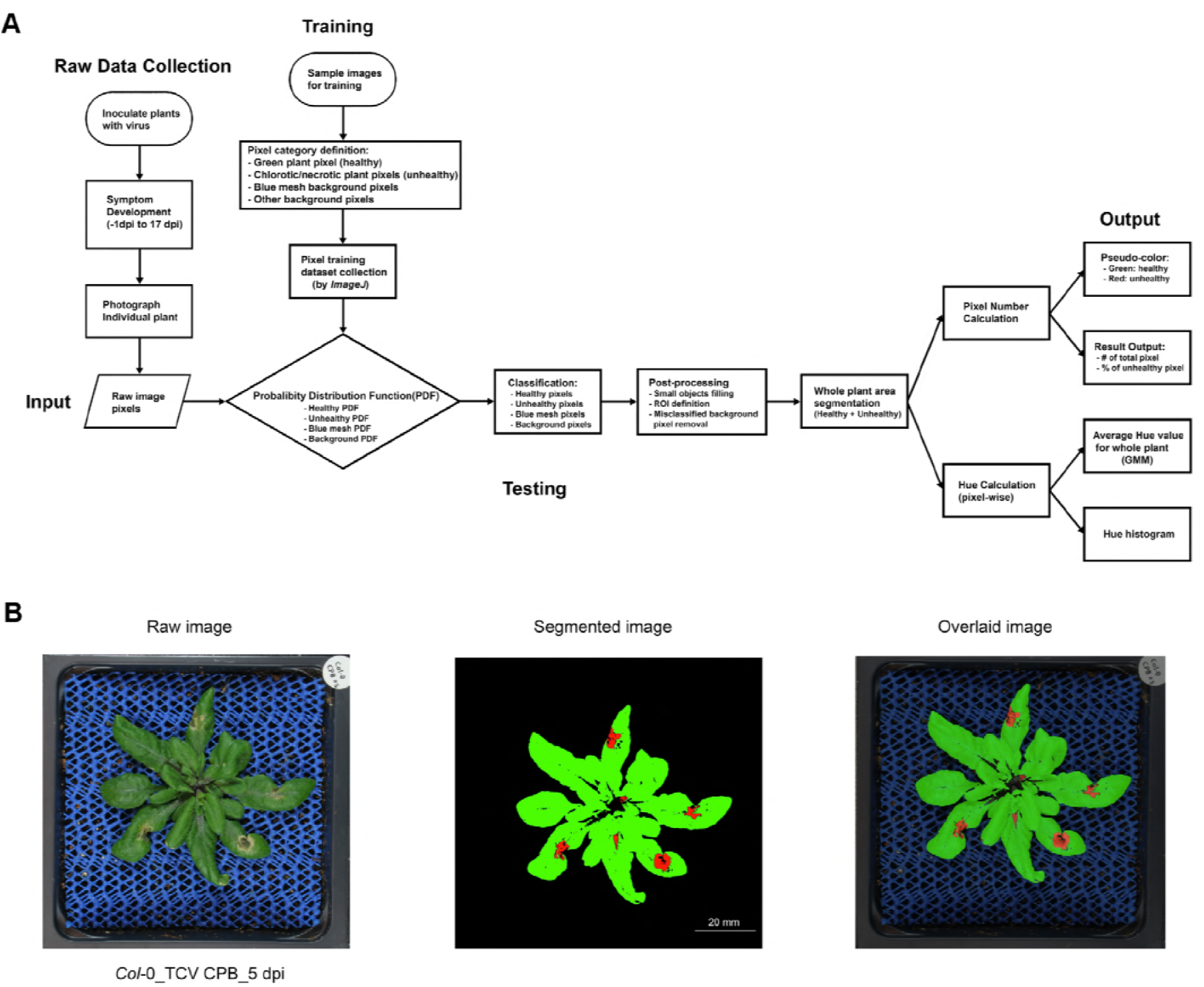
Image-based traits analysis workflow outline. (A) Workflow chart. Plants were inoculated with virus or mock solution. In the raw data collection section, Individual plant was photographed from 1 day before inoculation to 17 dpi. In the training session, representative sample images were chosen. Four pixel-classifiers were defined and RGB information of 1891 pixels in each class were collected to build up the training dataset. The Probability Distribution Function (PDF) for each category was calculated based on the training dataset. In the testing session, each pixel from an input image was calculated and classified into each class. After post-processing, the pipeline produced a summary of pixel number in each class and the hue value for each pixel in each raw image. Then appropriate statistics was applied to quantify and compare the output results among different groups. (B) One example of original raw plant image, segmented pseudo-color image, and overlaid image of *Col-0* plant inoculated with TCV CPB at 5 dpi.

To validate the approach, wild-type *(Col-0)* and hyper-susceptible *dcl2 dcl3 dcl4* mutant plants were inoculated with parental TCV or TCV CPB mutant virus, or were mock-inoculated (4 replicates per genotype and treatment combination). In the mock-inoculated groups, the majority of rosette area remained healthy (green) over time in both *Col-0* and *dcl2 dcl3 dcl4* plants (Figs 3A-B). Infection of both plant genotypes with parental TCV elicited local chlorosis at 7 dpi and nearly complete chlorosis/necrosis by 17 dpi (Fig 3A-B). Discoloration caused by parental TCV infection appeared to be more severe in the *dcl2 dcl3 dcl4* mutant plants than in wild-type controls (Fig 3B). TCV CPB inoculation led to parental virus-like discoloration symptoms in the *dcl* triple mutant plants (Fig 3B) but did not cause strong chlorosis symptoms in *Col-0* plants (Fig 3A). Although an increased number of chlorotic pixels was observed in *Col-0* rosettes at 15- and 17-days post TCV CPB inoculation, the increase was visually less than that caused by parental TCV (Fig 3A).

**Fig 3.**
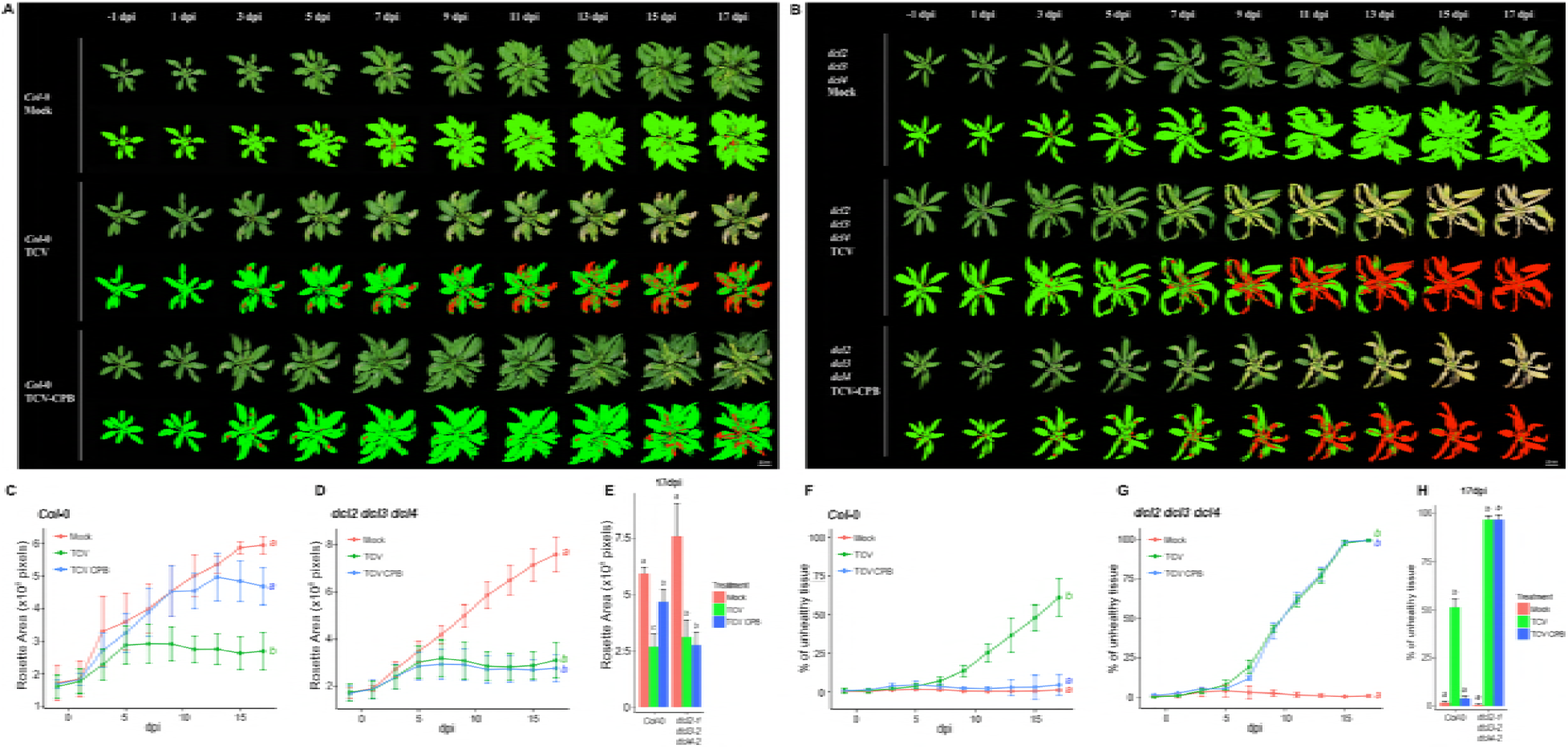
TCV infection-caused temporal changes in rosette size and the percentage of unhealthy tissues in control plants. (A) Temporal visualization of rosette leaves and the corresponding pseudocolor images of *Col-0* and (B) *dcl2 dcl3 dcl4* plants inoculated by TCV, TCV CPB inoculum or mock solution. Photos were taken from 1 day before inoculation to 17 dpi. In the pseudo-color images, green color referred to healthy plant pixels, while red color referred to chlorotic/necrotic (unhealthy) plant pixels. (C) The growth curves show the temporal change of rosette area (total pixel number, averaged by 4 plants, ± SE) in *Col-0* and (D) *dcl2 dcl3 dcl4* plants from 1 day before inoculation to 17 dpi (Kolmogorov-Smirnov test with α=0.05). (E) The bar-plot shows the average (+ SE, n=4) of rosette size at 17 dpi. Statistical analysis was calculated between treatments in each genotype. Bars with different letters are statistically different (Tukey *post hoc* test with α=0.05). (F) The curves show the percentage of unhealthy tissue change overtime (red pixels/total pixels, averaged by 4 plants, ± SE) in *Col-0* and (G) *dcl2 dcl3 dcl4* plants from 1 day before inoculation to 17 dpi (Kolmogorov-Smirnov test with α=0.05). (H) The bar-plot shows average (+ SE, n=4) of percentage of unhealthy pixels at 17 dpi. Statistical analysis was calculated between treatments in each genotype. Bars and curves with different letters are statistically different (Tukey *post hoc* test with α=0.05).

Parental TCV infection led to reduced rosette area, based on total number of pixels (healthy plus chlorotic), indicating a decrease in biomass. Parental TCV significantly inhibited rosette growth over time in both wild-type and hyper-susceptible controls (Figs 3C-D) (Kolmogorov-Smirnov test, p<0.05). Similarly, TCV CPB led to a comparable decrease in rosette area in *dcl2 dcl3 dcl4* mutants after 7 dpi, but not in *Col-0* (Figs 3B-D). At 17 dpi, a significant difference in rosette area was detected between the two virus-treatments in *Col-0,* but not in *dcl2 dcl3 dcl4* plants (Fig 3E).

The percentage of unhealthy chlorotic/necrotic pixels in the whole rosette was also calculated. Both parental TCV and TCV CPB infection caused a significant increase in the percentage of unhealthy tissues over time in *dcl2 dcl3 dcl4* plants (Fig 3G), from approximately 0% at 5 dpi to nearly 100% at 17 dpi (Fig 3G). In *Col-0* plants, parental TCV infection also led to a gradual increase in chlorotic tissue, from approximately 0% at 5 dpi to nearly 60% at 17 dpi (Fig 3F). In contrast, TCV CPB did not significantly change the percentage of chlorotic tissue in wild-type control plants over time (Fig 3F) (Kolmogorov-Smirnov test, p<0.05). At 17 dpi, no significant difference in the percentage of unhealthy tissues was detected in TCV CPB-inoculated and mock treatment *Col-0* plants (Fig 3H).

Measurement of hue value of leaf color has been used to estimate chlorophyll loss caused by biotic and abiotic stress [36–38]. Hue value information extracted from the RGB images was used as a parameter to quantify chlorosis in rosette leaves. The hue value quantifies color in terms of angle around a circle, with values ranging from 0° to 359° [39], starting in the red color range. Yellow and green hue ranges span from ~51° to 80° and ~81° to 140°, respectively. For each genotype and treatment combination, the mean proportion of pixels at each degree was plotted for each day. One primary peak centered around 100°, representing a green hue, was found in the histograms of mock-inoculated *Col-0* and *dcl2 dcl3 dcl4* plants over time (left panel, Figs 4A-B). After 7 dpi, parental TCV infection caused a shift from a unimodal distribution of green hue values to a bimodal distribution that included a peak of yellow hue values in both genotypes (middle panel, Figs 4A-B). Notably, this green to yellow shift observed in *dcl2 dcl3 dcl4* mutants was more complete than that in *Col-0* plants at 17 dpi, suggesting the hyper-susceptible control leaves were more yellow than wild-type leaves (middle panel, Figs 4A-B). Similar to parental virus, TCV CPB induced a gradual shift from green to yellow in *dcl2 dcl3 dcl4* plants (right panel, Fig 4B). In contrast, this green-to-yellow shift was not observed in *Col-0* plants inoculated with TCV CPB (right panel, Fig 4A).

**Fig 4.**
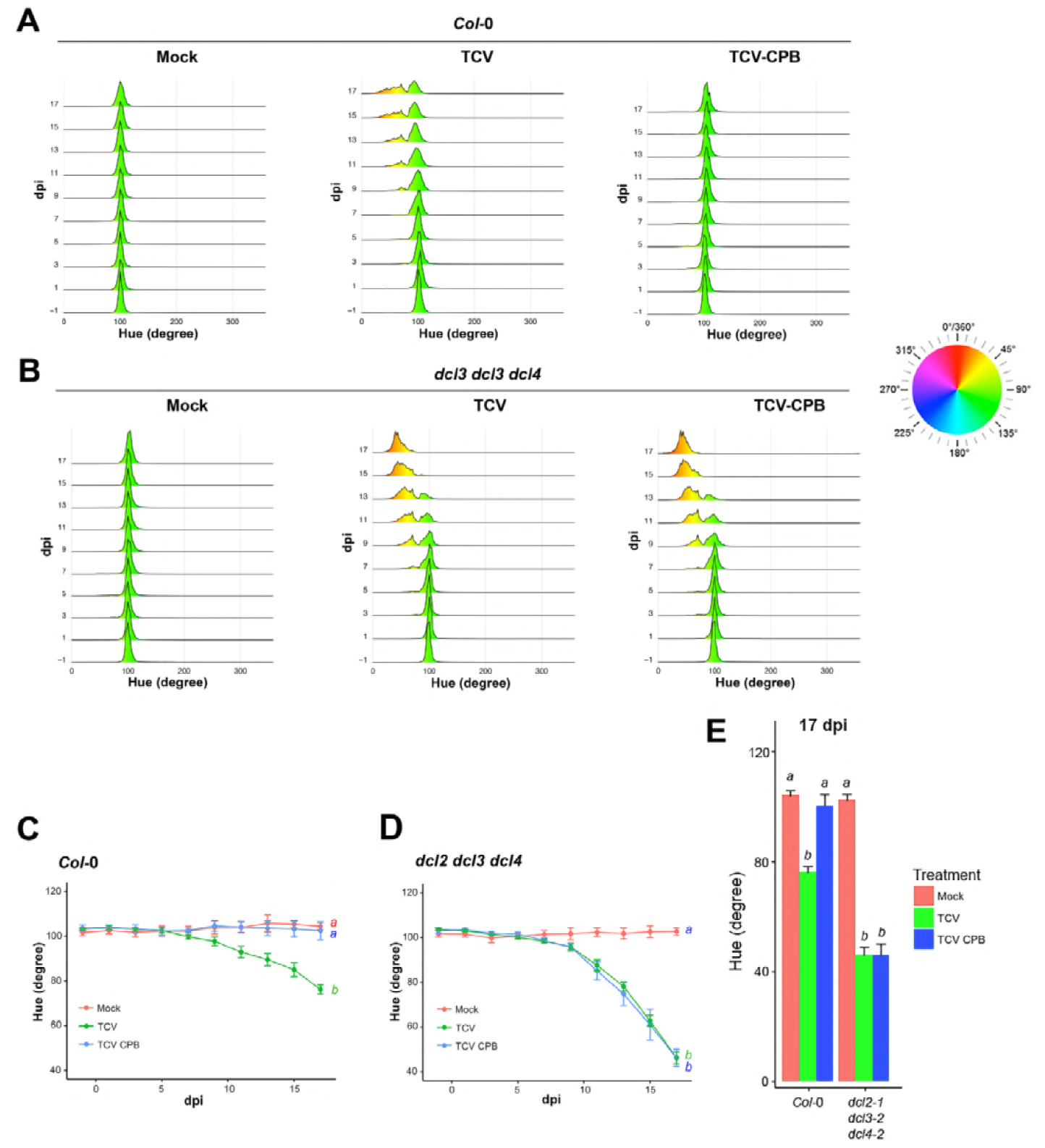
TCV infection-caused temporal color changes in control plants. (A) The histogram illustrates the temporal changes (1 day before inoculation to 17 dpi, from bottom to above) of pixel distribution at each hue value (degree) in *Col-0* and (B) *dcl2 dcl3 dcl4* plant rosette images. Pixels were collected and summed up from four individual images in the same treatment group: mock (left panel), TCV (middle panel), or TCV CPB (right panel) inoculum. The color chart illustrates the relationship between the hue value (degree) with the corresponding RGB coordinates. (C) The plots show the average hue value (degree) change over time (averaged by 4 plants, ± SE) in *Col-0* and (D) *dcl2 dcl3 dcl4* plants from 1 day before inoculation to 17 dpi (Kolmogorov-Smirnov test with α=0.05). (E) The bar plot shows the average hue value (+ SE, n=4) at 17 dpi in *Col-0* and *dcl2 dcl3 dcl4* plants. Statistical analysis was calculated between treatments in each genotype. Bars and curves with different letters are statistically different (Tukey *post hoc* test with α=0.05).

The average hue value for whole plants in each treatment group was calculated. Parental TCV infection caused a temporal decrease in hue value in both *Col-0* (from 100° to 75°) and *dcl2 dcl3 dcl4* plants (from 100° to 40°) (Figs 4C-D). TCV CPB induced a similar decrease in average hue value over time in *dcl2 dcl3 dcl4* mutants, but did not significantly affect hue value in *Col-0* plants (Figs 4C-E). These results were consistent with the healthy/chlorotic classification-based results (Fig 3). These data indicate that the image-based growth and color trait measurement protocols were effective in quantifying virus-induced symptoms in *Arabidopsis,* and in distinguishing plant susceptibility or virus virulence phenotypes.

### Image-based analysis of TCV and TCV CPB infection in *ago* mutants

As with the *dcl2 dcl3 dcl4* plants, we hypothesized that loss of *ago* genes with a function in anti-TCV silencing would be revealed by gain of susceptibility to the VSR-deficient TCV CPB mutant. A set of ten mutant plants with defects in each of the ten *Arabidopsis AGO* genes was inoculated with mock solution, parental TCV or TCV CPB, and image-based traits and virus protein levels over an infection time-course were measured. Pairwise Kolmogorov-Smirnov (K-S) tests were done to compare time-series data.

Parental TCV infection caused a significant decrease in rosette size over time in all genotypes (Fig 5A). In contrast, TCV CPB inoculation led to three different effects, depending on genotype : a) no significant effect on rosette size overtime in *ago4-2, ago5-2, ago8-1, ago9-5* and *ago10-5*; b) significant negative effects on rosette size change over time in *ago2-1, ago3-2, zip-1 (ago7* mutant) and *dcl2 dcl3 dcl4* control; or c) observable, but not significant, negative effects on rosette size change over time in *ago1-27, ago6-3,* and *Col-0* plants (Fig 5A). After 7 dpi, a significant increase in the percentage of chlorotic/necrotic tissue was observed in all genotypes infected with parental TCV (Fig 5B), though the degree of increase varied in different genotypes (Fig 6C). In the *dcl2 dcl3 dcl4* mutant, both parental and mutant TCV caused a comparable increase in the percentage of unhealthy tissue (Fig 5B). Among the ten *ago* mutants, TCV CPB significantly increased the percentage of unhealthy tissue only in *ago2-1* and *zip-1* mutants over time (Fig 5B). The *ago1-27, ago3-2, ago4-2, ago5-2, ago6-3, ago8-1, ago9-5* and *ago10-5* mutants were not responsive to TCV CPB inoculation (Fig 5B). Focusing on the data on 17 dpi, the effects caused by TCV CPB on rosette chlorosis in *ago2-1* and *zip-1* mutant could be directly displayed (Fig 6A) and statistically tested (Fig 6C) (Tukey *post hoc* test with α=0.05). Similarly, by using this statistic test, we further confirmed that TCV CPB caused a significant decrease in *ago2-1, ago3-2* and *zip-1* rosette size at 17 dpi (Fig 6B).

**Fig 5.**
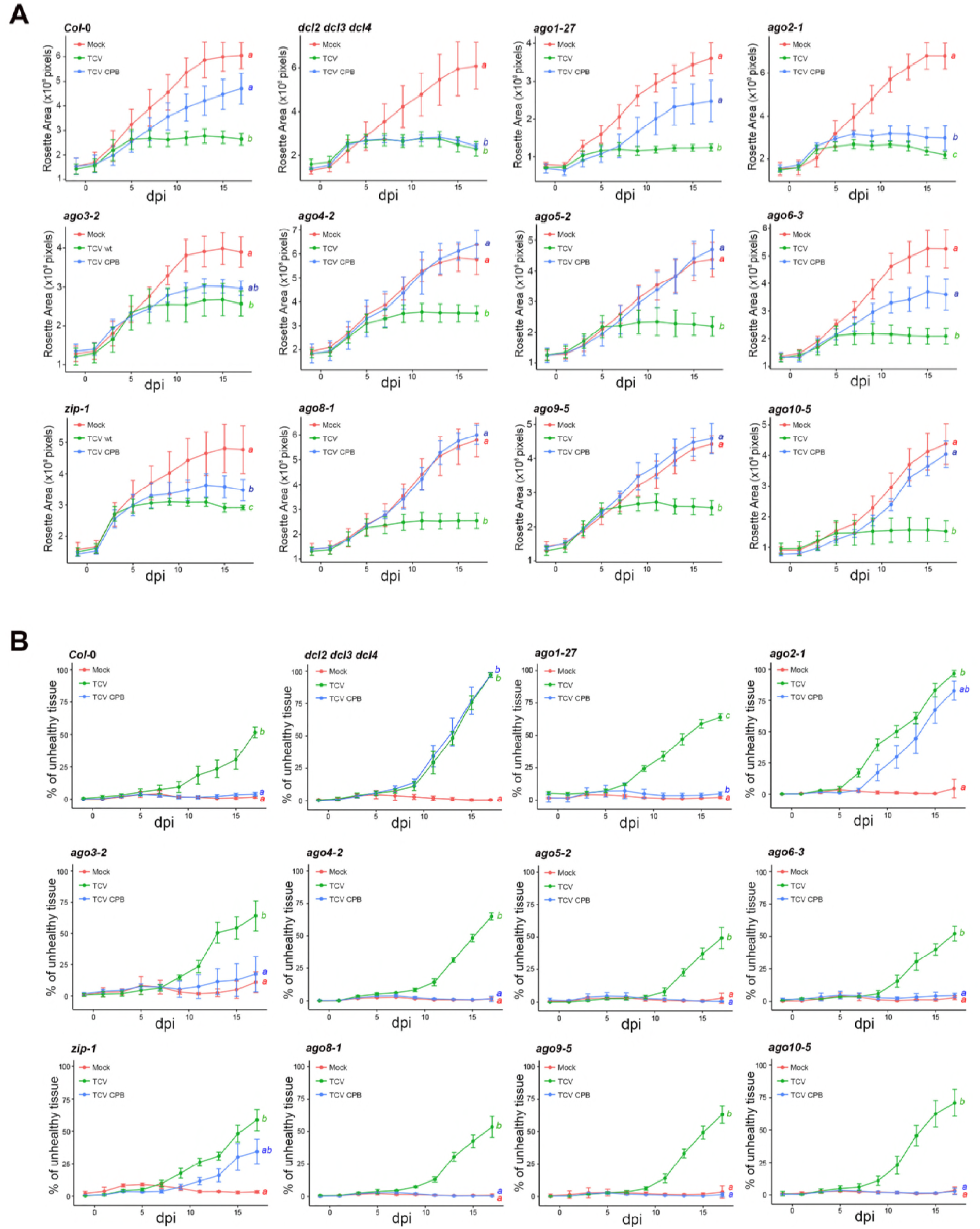
TCV infection-caused temporal changes in rosette size and the percentage of unhealthy tissues in single *ago* mutant plants. (A) The growth curves show the temporal change of rosette area (total pixel number, averaged by 4 plants, ± SE) in *Col-0, dcl2 dcl3 dcl4* and ten single *ago* mutant plants from 1 day before inoculation to 17 dpi (Kolmogorov-Smirnov test with α=0.05, detailed D value and p value are listed in S1 Table). (B) The curves show the percentage of unhealthy tissue change overtime (red pixels/total pixels, averaged by 4 plants, ± SE) in *Col-0, dcl2 dcl3 dcl4* and ten single *ago* mutant plants from 1 day before inoculation to 17 dpi. (Kolmogorov-Smirnov test with α=0.05, detailed D value and p value are listed in S2 Table). Curves with different letters are statistically different.

**Fig 6.**
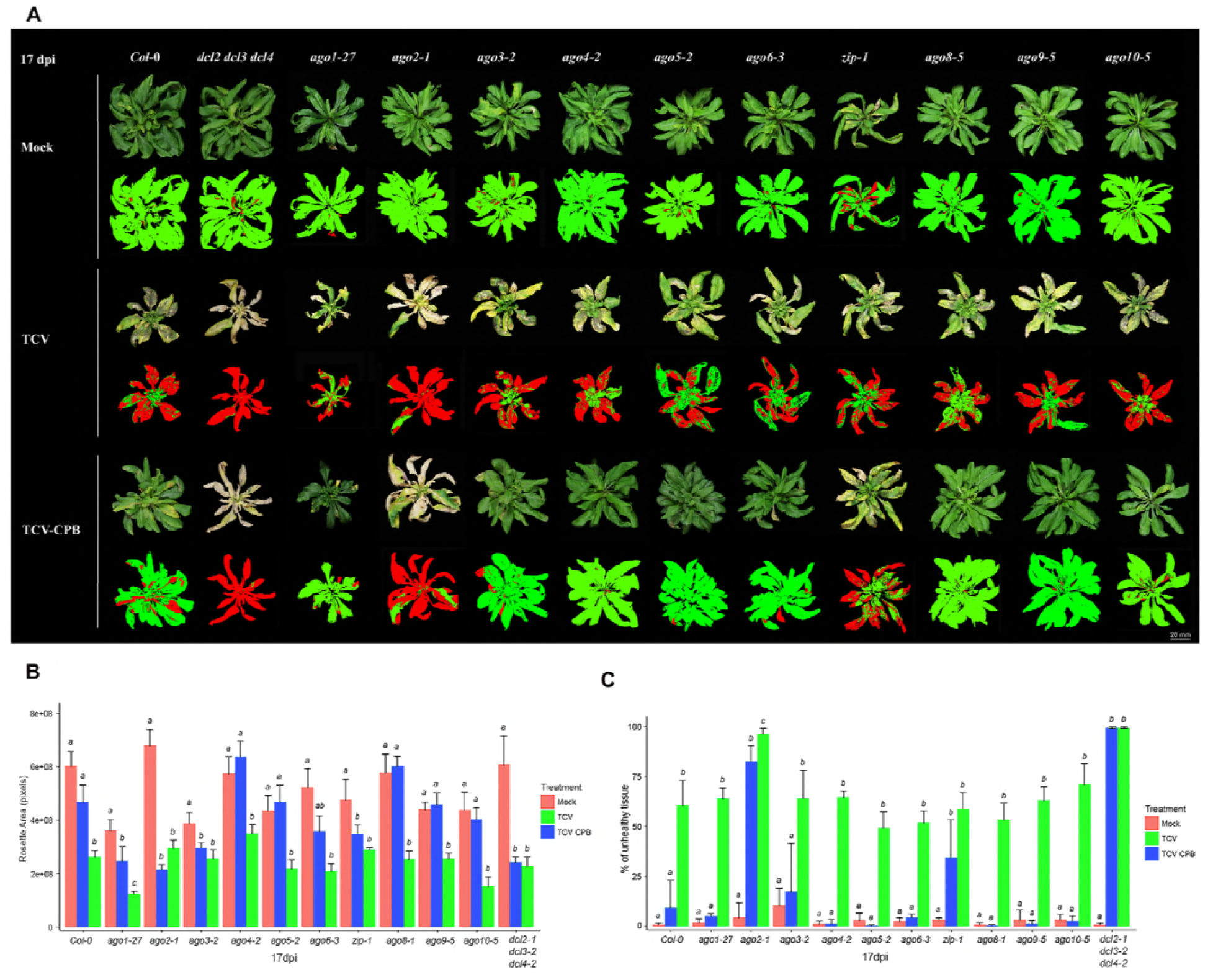
Rosette size and the percentage of unhealthy tissue in single *ago* mutant plants infected with TCV at 17 dpi. (A) Representative visualization of rosette leaves and the corresponding pseudocolor images in *Col-0, dcl2 dcl3 dcl4* and ten single *ago* mutant plants, which were separately inoculated by mock solution, TCV or TCV CPB inoculum. Photos were taken at 17 dpi. In the pseudo-color images, green color referred to healthy plant pixels, while red color referred to chlorotic/necrotic (unhealthy) plant pixels. (B) The bar plot shows the average rosette area (total pixel number, n=4, + SE) in *Col-0, dcl2 dcl3 dcl4* and ten single *ago* mutant plants at 17 dpi. (C) The bar plot shows the average percentage of unhealthy tissue change overtime (red pixels/total pixels, n=4, + SE) in *Col-0, dcl2 dcl3 dcl4* and single *ago* mutant plants at 17 dpi. Statistical analysis was calculated between treatments in each genotype. Bars with different letters are statistically different (Tukey *post hoc* test with α=0.05).

Another output of the phenotyping pipeline to evaluate leaf color is hue value (Fig 2A). The temporal changes in hue value in the ten inoculated *ago* mutants and control plants were measured over time (Fig 7). One primary peak at approximately 100° (green) was observed in early stages for all genotype and inoculation combinations (−1 to 5 dpi in each panel, Fig 7). In the mock-inoculated group, the primary peak remained at around 100° in all genotypes throughout the time course. The green peak gradually shifted to yellow in each *ago* mutant and the two controls infected with parental TCV (7 to 17 dpi in each panel, Fig 7). As expected, *dcl2 dcl3 dcl4* mutant plants infected with TCV CPB turned from green (peak at 100°) to yellow over time (blue area, dcl2/3/4 panel, Fig 7). Similarly, a complete green-to-yellow shift was measured in *ago2-1* mutant plants infected with TCV CPB (blue area in *ago2-1* panel, Fig 7). Gradual, partial shift from green to yellow was measured in *zip-1* plants infected with TCV CPB (blue area in *zip-1* panel, Fig 7). All other *ago* mutants *(ago1-27, ago3-2, ago4-2, ago5-2, ago6-3, ago8-1, ago9-5* and *ago10-5*) inoculated with TCV CPB were similar to mock-inoculated plants (blue area, Fig 7).

**Fig 7.**
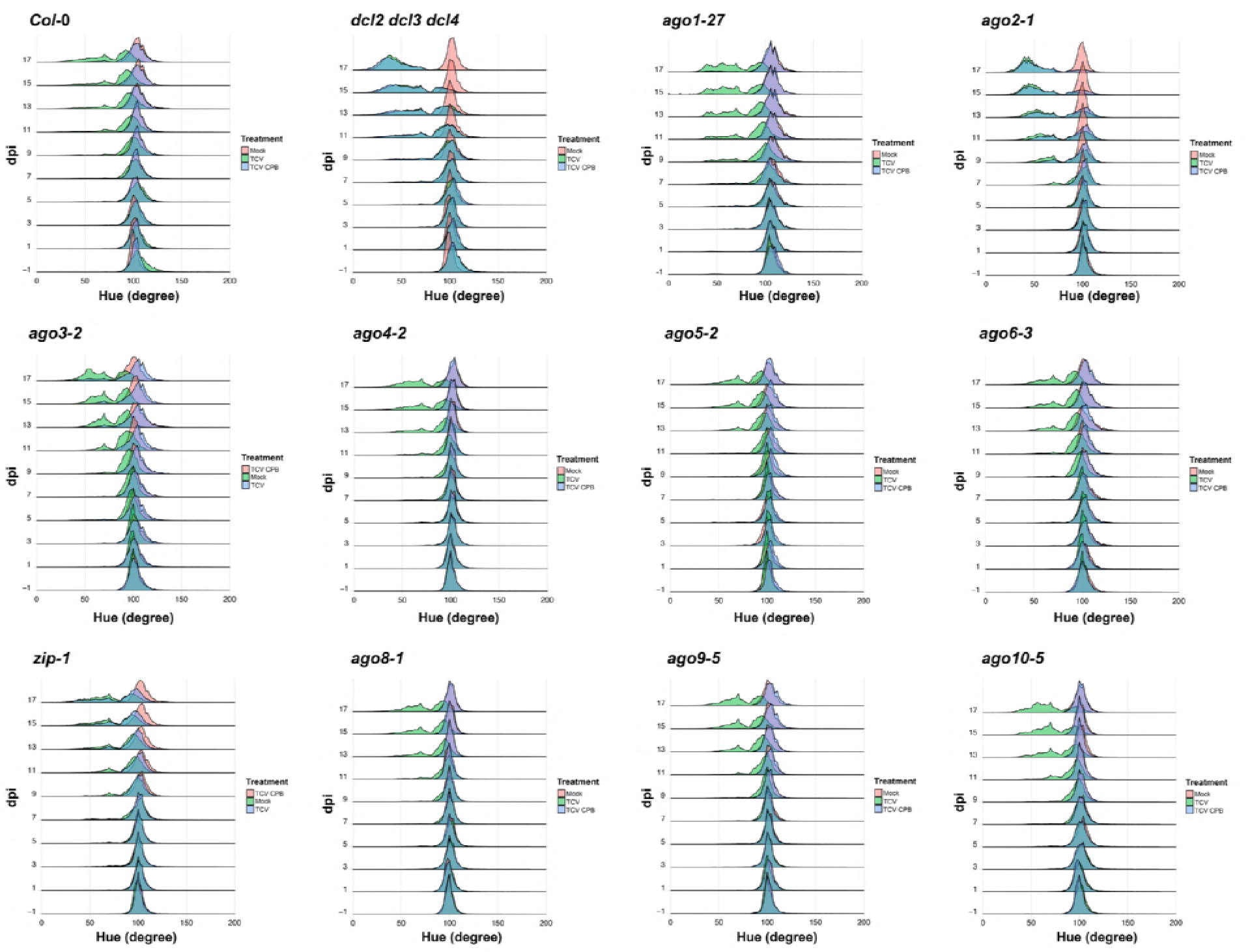
TCV infection-caused temporal pixel distribution changes in single *ago* mutant plants. Each histogram illustrates the temporal changes (1 day before inoculation to 17 dpi, from bottom to above) of pixel distribution at each hue value (degree) in *Col-0, dcl2 dcl3 dcl4* or the ten single *ago* mutant rosette images. Pixels were collected and summed up from four individual images in the same treatment group. For the separated distribution histogram in each genotype and treatment combination, refers to S12 Fig.

To quantify the color shift observed above, average hue value was calculated. Pairwise Kolmogorov-Smirnov (K-S) tests were applied to compare time-series data. Parental TCV infection led to a gradual decrease in average hue value in all genotypes (Fig 8A), while TCV CPB caused a similar decrease only in *ago2-1* and *zip-1* mutants (Fig 8A) In addition, data at 17 dpi further confirmed the negative effects of TCV CPB on average hue value in *ago2-1* and *zip-1* mutants (Fig 8B) (Tukey *post hoc* test with α=0.05).

**Fig 8.**
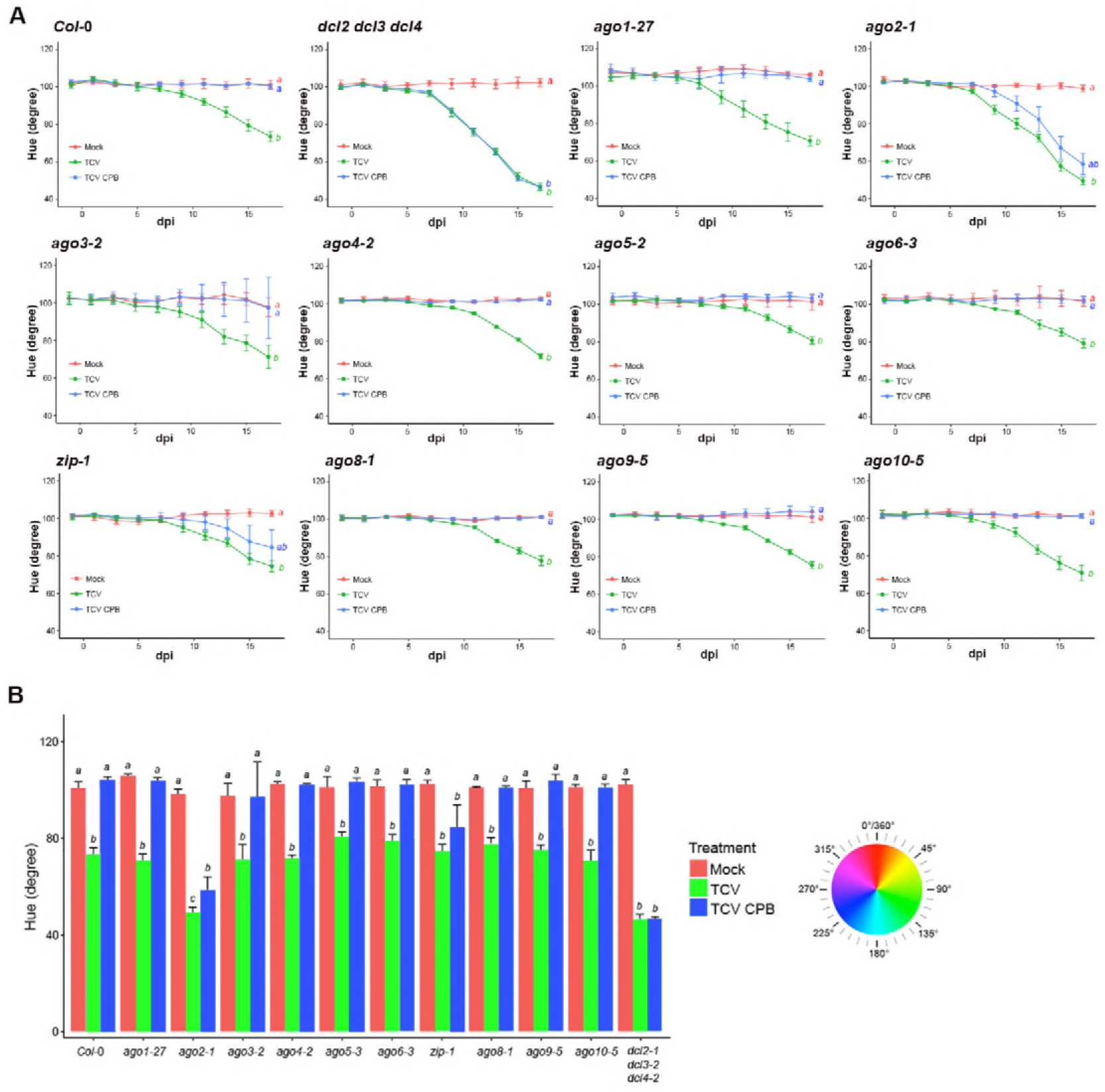
TCV infection-caused temporal average hue value changes in single *ago* mutant plants. (A) The plots show the average hue value (degree) change over time (averaged by 4 images, ± SE) in *Col-0, dcl2 dcl3 dcl4* and single *ago* mutant plants from 1 day before inoculation to 17 dpi (Kolmogorov-Smirnov test with α=0.05, detailed D value and p value are listed in S3 Table). (B) The bar-plot shows the average hue value (+ SE, n=4) in *Col-0, dcl2 dcl3 dcl4* and single *ago* mutant plants at 17 dpi. Statistical analysis was calculated between treatments in each genotype. Bars and curves with different letters are statistically different (Tukey *post hoc* test with α=0.05). The color chart illustrates the relationship between the hue value (degree) with the corresponding RGB coordinates.

### Analysis of viral coat protein levels in *ago* mutants

To further investigate the roles of anti-TCV AGO proteins identified above in different tissues, immunoblotting assay was used to analyze viral coat protein (P38) levels in inoculated, non-inoculated (systemic) tissues, and inflorescence tissues of *ago* mutants and control plants. High levels of P38 were detected in leaves and inflorescence tissues of all ten single *ago* mutants and two control plants inoculated with parental TCV (left panel, Fig 9A; blue bars in Figs 9B-D). Local accumulation of CPB-P38 was only detected in *ago1-27, ago2-1,* and *zip-1* mutant rosette leaves (right panel, Fig 9A, red bars in Fig 9B), at a comparable level with that in *dcl2 dcl3 dcl4* leaves (red bars in Fig 9B). In non-inoculated cauline leaves, systemic accumulation of CPB-P38 was only detected in *ago2-1* and *zip-1* mutants (Fig 9C), and at levels that were approximately half of that in *dcl2 dcl3 dcl4* mutants (Fig 9C). CPB-P38 in local and systemic leaves was also detected in two of four biological replicates of *ago3-2* mutants (right panel, Fig 9A), though at a significantly lower level than that in *ago2-1* mutants (Figs 9B-C). Notably, CPB-P38 was not detected in inflorescence tissues of any of the ten *ago* single mutant and *Col-0* plants (right panel, Figs 9A and 9C).

**Fig 9.**
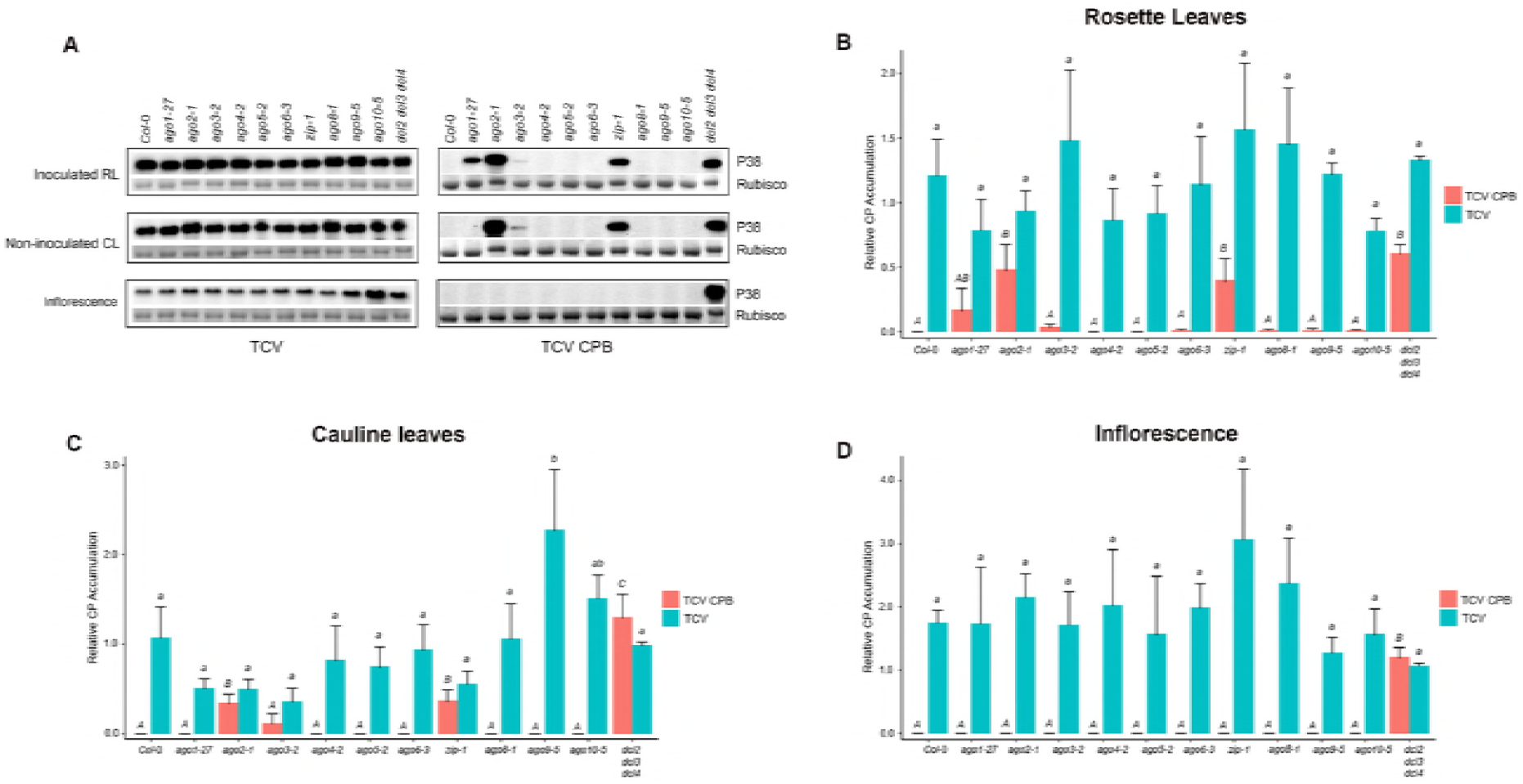
Local and systemic accumulation of TCV CP in single *ago* mutant plants. (A) TCV viral coat protein (P38) accumulation in different tissues of *Col-0, dcl2 dcl3 dcl4* and ten single *ago* mutant plants infected with TCV (left panel) or TCV CPB (right panel). RL: rosette leaves; CL: cauline leaves; Inflorescence. Rubisco protein: internal control. (B) Summary data: the local level of parental or CPB mutant P38 accumulation in inoculated rosette leaves of *Col-0, dcl2 dcl3 dcl4* and the ten single *ago* mutant plants at 7 dpi. (C) Summary data: the systemic level of parental or CPB mutant P38 accumulation in non-inoculated cauline leaves of *Col-0, dcl2 dcl3 dcl4* and the ten single *ago* mutant plants at 14 dpi. (D) Summary data: the systemic level of parental or CPB mutant P38 accumulation in inflorescence tissues of *Col-0, dcl2 dcl3 dcl4* and the ten single *ago* mutant plants at 14 dpi. All bar plot shows average (+SE) of four biological replicates, expressed relative to *dcl2 dcl3 dcl4.* Bars with different letters are statistically different (Tukey *post hoc* test with α=0.05).

Next, to investigate if AGO2 and AGO7 had additive antiviral effects, and also to test if the antiviral activities of AGO1, the previously reported antiviral AGO protein against TCV [14], was masked by AGO2 or AGO7, we performed immunoblotting experiments as described above in single, double or triple mutants combining *ago1-27, ago2-1* and/or *zip-1* mutant allele (Fig 10). First, parental TCV CPs accumulated at comparable levels in inoculated rosette, non-inoculated cauline leaves and inflorescence tissue in the single, double and triple *ago* mutants (Figs 10A-C). In local and systemic leaves, no significant differences in P38 level were observed between *ago2-1, zip-1* and *ago2-1 zip-1* double mutant (Figs 10B-C). These results suggested that AGO2 and AGO7 play non-additive antiviral roles in *Arabidopsis* leaves during TCV infection. To examine if AGO1, AGO2 or AGO7 play redundant antiviral roles, we checked if down-regulating *ago1* had any enhancing effects on P38 accumulation in *ago2* or *ago7* mutants. We found no significant differences in local P38 level in rosette leaves between *ago2-1* single, *zip-1* single, *ago1-27 ago2-1* double, *ago1-27 zip-1* double and *ago1-27, ago2-1 zip-1* triple mutant (Fig 10B). Similarly, combining the *ago1-27* allele with an *ago2-1* or *zip-1* allele, or *ago2-1* and *zip-1,* did not significantly affect the CPB-P38 accumulation level in systemic cauline leaves (Fig 10C). These results suggested that introducing the *ago1-27* allele does not enhance or suppress the antiviral activities of AGO2 and AGO7 in *Arabidopsis* leaves. Similar to the observation in *ago* single mutants, no TCV CPB CPs were detected in the inflorescence clusters of the double and triple mutants tested (Fig 10D).

**Fig 10.**
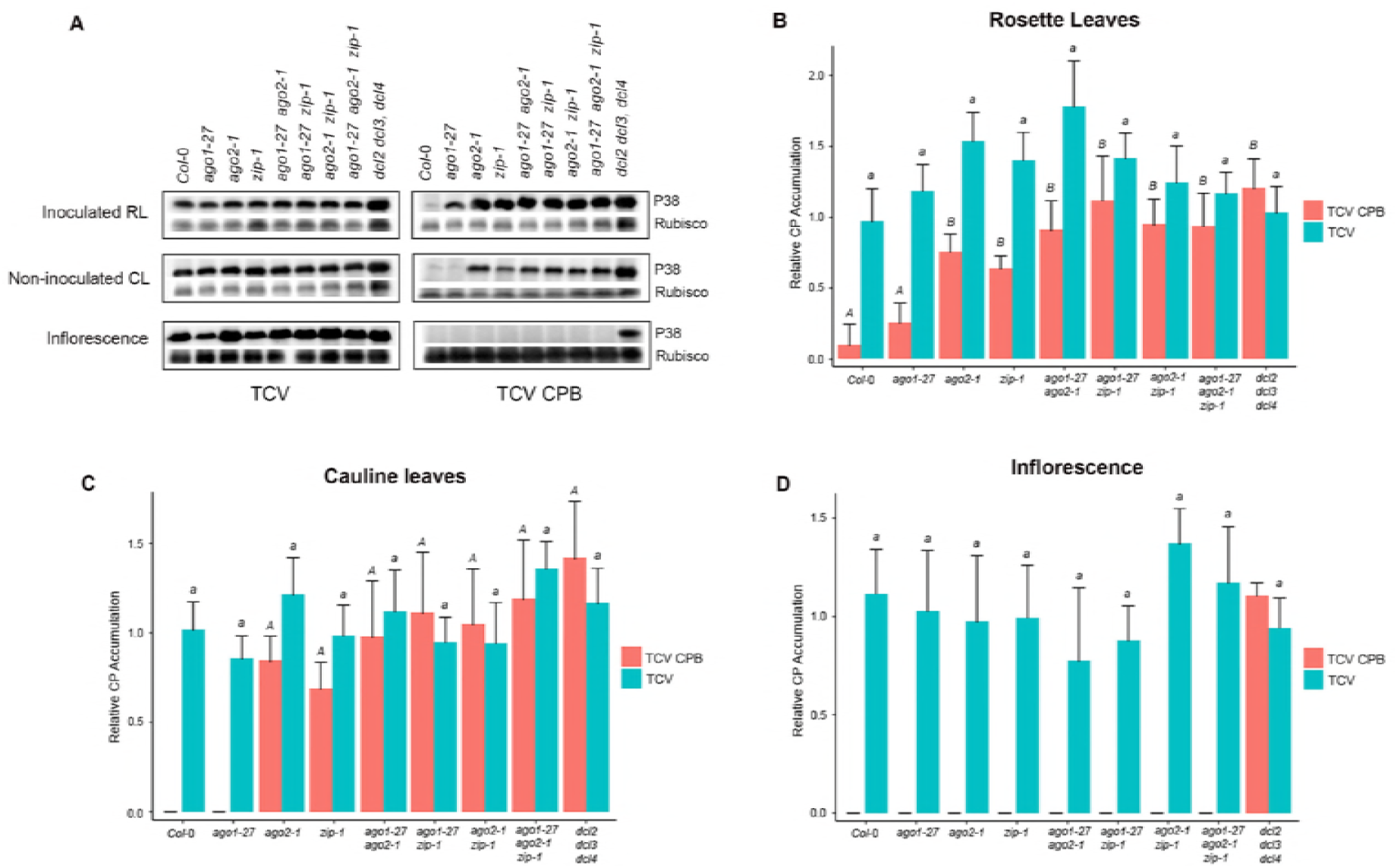
Local and systemic accumulation of P38 in a selected group of double and triple *ago* mutant plants. (A) TCV viral coat protein (P38) accumulation in different tissues of *Col-0, dcl2 dcl3 dcl4* and *ago1-27, ago2-1, zip-1* single, double or triple mutant plants infected with TCV (left panel) or TCV CPB (right panel). RL: rosette leaves; CL: cauline leaves; Inflorescence. Rubisco protein: internal control. (B) Summary data: the local level of parental or CPB mutant P38 accumulation in inoculated rosette leaves of *Col-0, dcl2 dcl3 dcl4* and *tago1-27, ago2-1, zip-1* single, double or triple mutant plants at 7 dpi. (C) Summary data: the systemic level of parental or CPB mutant P38 accumulation in non-inoculated cauline leaves of *Col-0, dcl2 dcl3 dcl4* and *ago1-27, ago2-1, zip-1* single, double or triple mutant plants at 14 dpi. (D) Summary data: the systemic level of parental or CPB mutant P38 accumulation in inflorescence tissues of *Col-0, dcl2 dcl3 dcl4* and *ago1-27, ago2-1, zip-1* single, double or triple mutant plants at 14 dpi. All bar plot shows average (+SE) of four biological replicates, expressed relative to *dcl2 dcl3 dcl4.* Bars with different letters are statistically different (Tukey *post hoc* test with α=0.05).

### Analysis of plant height during TCV infection in *ago* mutants

The effects of TCV infection on plant height in ten *ago* mutants, the *dcl* triple mutant and wild-type plants inoculated with parental and VSR-deficient TCV were measured. Growth curves were plotted from one day pre-inoculation to 21 dpi as described previously [40]. Pairwise Kolmogorov-Smirnov (K-S) tests were done to compare time-series data.

Under mock treatment, plant height was greater than 30 cm in most genotypes at 21 dpi, except in *ago1-27* and *dcl2 dcl3 dcl4* mutants (Figs 11A and 11C). TCV infection led to reduced plant height (<10 cm) starting around 7 dpi in all genotypes (Fig 11A). The TCV CPB mutant virus had only minor effects on height of inoculated *Col-0* plants over time, but strongly affected height in *dcl2 dcl3 dcl4* mutant plants (Figs 11A-C). Mild inhibition of height caused by TCV CPB was observed at intermediate time points *(Tukey post hoc* test, α=0.05, Fig 11B), though no effects on plant height were detected at 21 dpi (Fig 11C).

**Fig 11.**
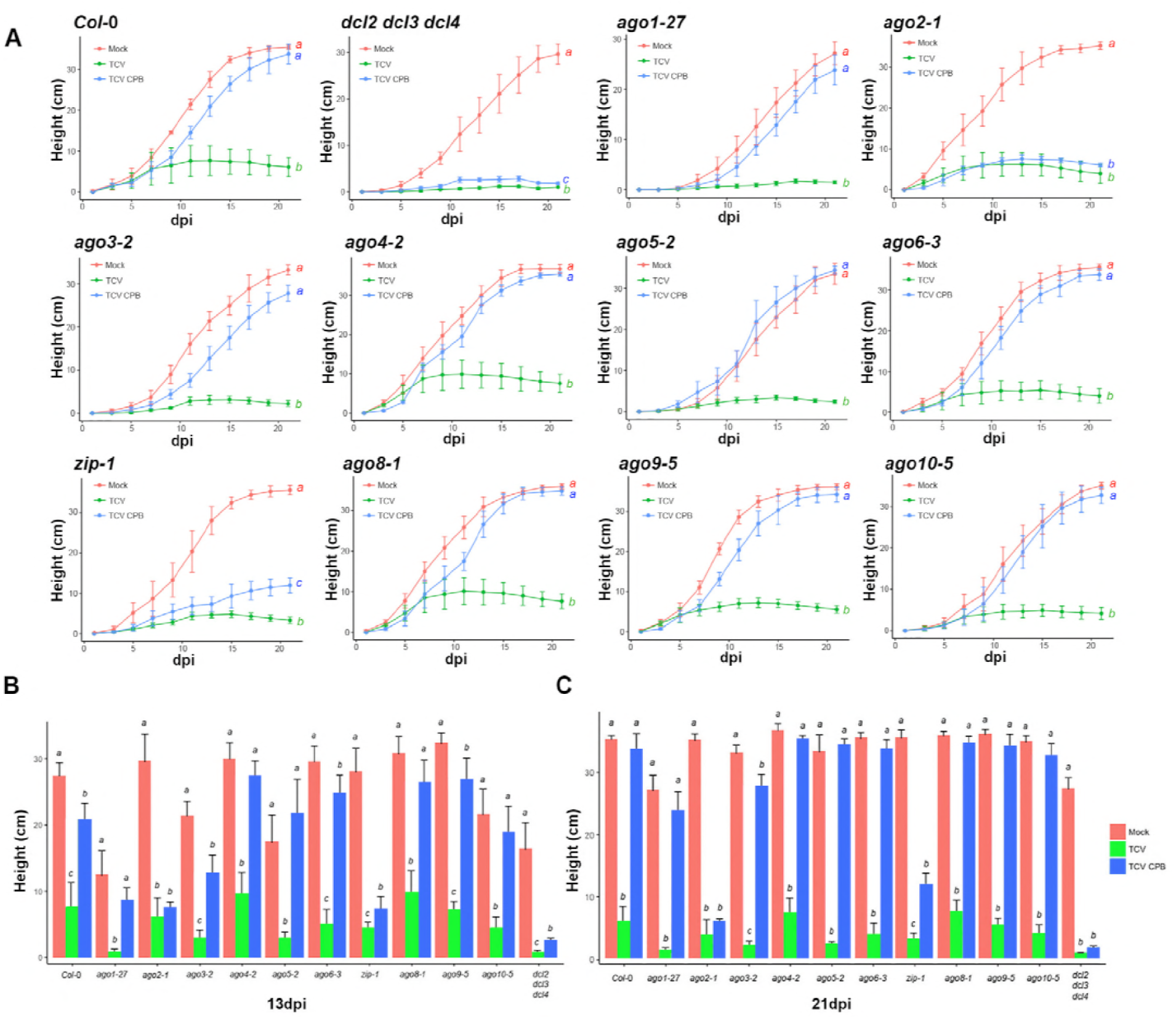
The effects of TCV infection on plant growth in single *ago* mutant plants. (A) The histograms show the plant growth curve (averaged by 6 plants, ± SE) in *Col-0, dcl2 dcl3 dcl4* and ten single *ago* mutant plants inoculated with mock solution, TCV or TCV CPB inoculum from one day before inoculation day to 21 dpi (Kolmogorov-Smirnov test with α=0.05, detailed D value and p value are listed in S4 Table). (B) The bar plot shows the average plant height (+ SE, n=6) in *Col-0, dcl2 dcl3 dcl4* and ten single *ago* mutant plants at 13 dpi or (C) 21 dpi. Statistical analysis was calculated between treatments in each genotype. Bars and curves with different letters are statistically different (Tukey *post hoc* test with α=0.05).

The TCV CPB mutant also strongly inhibited plant height in *ago2-1* and *zip-1* mutants (Fig 11A). In *ago2-1* mutants, TCV CPB infection had parental virus-like effects on plant height at 13 and 21 dpi (Figs 11 B-C). In *zip-1* mutants, the suppressor-deficient virus led to an approximately 60% decrease in plant height compared with mock-inoculated control plants at both time points (Figs 11B-C). In *ago3-2* mutant plants, TCV CPB caused modest reduction of reduced height at both 13 dpi and 21 dpi (by *Tukey post hoc* test with α=0.05, Fig 11C). TCV CPB had little or no effects on plant height in *ago1-27, ago4-2, ago5-2, ago6-3, ago8-1, ago9-5,* or *ago10-5* mutants.

## Discussion

Precisely measuring biotic and abiotic stress traits is important to understand the impact of the environment, pathogens and pests on plant growth and development, and to assist breeders to improve crops in modern agriculture. Objective, reproducible, and accurate quantification of disease severity is important for development and testing of disease-resistant crops. Various methods to estimate disease severity in plants are in use [30, 41–43], though many of these are dependent on qualitative or subjective observations, or require destructive sampling of plant material. Recently, some non-destructive, image-based analysis methods have been developed to measure plant morphology [44–46]. Machine learning techniques have been applied to efficiently process color trait data from large numbers of images, and to link the inputs to the outputs mathematically [47, 48]. Here, we integrated a machine learning algorithm and analysis tools in the open-source phenotyping platform PlantCV [32, 33] to develop an image-based disease trait analysis pipeline to objectively measure disease severity in *Arabidopsis* plants infected by viruses.

In this pipeline, the naïve Bayes machine learning method [31, 32] was used for segmenting plant tissue from background in an image at pixel-level. First, four naïve Bayes classifiers were defined: 1) green plant pixels (“healthy”); 2) chlorotic/necrotic plant pixels (“unhealthy”); 3) blue mesh background pixels; 4) other background pixels. Since the light intensity condition was relatively constant, the color information of pixels from small number of sample images was sufficient for generating the training dataset to cover the range of variation of all images. In the training session, four classes of pixels could be segmented simultaneously. In addition, the naïve Bayes segmentation process is robust across large number of images due to its probabilistic nature. This pipeline is also simple and computationally expedient. The output of the pipeline could be statistically analyzed to quantify plant size, the proportion of unhealthy tissue, and leaf color. This non-destructive phenotyping pipeline enables visualization and quantification of disease symptom development over time from large numbers of plants.

In this study, the image-based disease analysis method was put to the test in wildtype and mutant Arabidopsis plants, with defects in *AGO* genes, infected by parental and VSR-defective TCV. Detailed phenotyping of inoculated Arabidopsis *ago* mutant plants was done. Ten AGO genes are encoded in the *Arabidopsis* genome and they are grouped into three major clades: AGO1/5/10, AGO2/3/7, and AGO4/6/8/9 [3]. Combining disease phenotyping and biochemical results, AGO2 and AGO7 were identified as prominent factors during anti-TCV defense, along with minor antiviral roles of AGO3. The results with AGO2 and AGO7 are consistent with previous studies [14, 20]. Using the VSR-defective virus, the antiviral role of AGO7 was relatively minor compared to that of AGO2. AGO2 and AGO7 were non-additive in leaves (Fig. 10). However, movement of the TCV mutant to inflorescence tissues was still inhibited in *ago2 ago7* double mutant, but not in *dcl2 dcl3 dcl4* mutants (Fig. 10). These results implied that other AGOs not tested in the current genetic combinations are involved to restrict TCV movement to, or accumulation in, inflorescence tissues.

AGO1 was previously considered as a prominent antiviral factor against several viruses [14, 17, 18]. However, we observed that the *ago1-27* mutant was not susceptible to the VSR-defective TCV, indicating that full AGO1 activity is clearly not necessary for anti-TCV function. Since AGO1 is critical for regulating gene expression in numerous developmental and physiological pathways [3], *ago1* null alleles are lethal and difficult to test. Thus, in the hypomorphic *ago1-27* mutant, either enough functional AGO1 protein is present to contribute to suppression of the mutant TCV, or there is little or no role for AGO1 in the anti-TCV response. It should be noted that disruption of full AGO1 function can up-regulate of AGO2 expression [17, 20], meaning that AGO2 levels in the *ago1-27* mutant could affect antiviral function of AGO2. Therefore, our data do not exclude possible direct or indirect roles for AGO1 during TCV infection.

Our finding that *ago2* mutants were more susceptible to the VSR-defective TCV provided additional evidence that AGO2 functions as an important antiviral effector against a broad spectrum of plant viruses [15, 20–22]. We also found minor effects of AGO3. A recent report proposed that AGO3 was not involved in the antiviral pathway against *cucumber mosaic virus* (CMV) [49], though this conclusion is complicated by the use of parental CMV containing a functional VSR, which could possibly mask the effects of AGO3. *AGO3* and *AGO2* belongs to the same phylogenetic clade, and these two genes are closely linked in the *Arabidopsis* genome. AGO3 has slicer activity against viral RNAs *in vitro* [50] and binds siRNAs derived from *Potato spindle tuber viroid in vivo* [51]. Therefore, some degree of antiviral activity of AGO3 against TCV, as revealed using the VSR-defective virus, might not be surprising. However, to further investigation with combinations of mutations, such as with an *ago2 ago3* double mutant, would be informative.

AGO7 binds miR390, which targets TAS3 transcripts for biogenesis of tasiRNAs [52-55]. AGO7 has also been shown to affect antiviral function [14, 15], though its role is not clear. AGO7 has slicer activity [56], but it is not clear if AGO7 directly interacts with viral siRNAs or genomic (or subgenomic) RNAs during infection.

Our findings partly agreed with a previously reported modular, tissue-specific mode of antiviral AGO function against *Turnip mosaic virus* (TuMV), with AGO2 possessing a major role and AGO10, AGO5 and AGO7 having minor effects [15]. With TCV, both AGO2 and AGO7 are influential in protecting leaves, along with minor contributions from AGO3. This may reflect the fact that the VSRs encoded by different viruses use different molecular strategies to affect antiviral effector functions [57]. HC-Pro from TuMV was found to sequester viral siRNAs away from multiple AGO proteins [15]. Previous reports suggested that TCV infection led to expression changes of many endogenous genes, including AGO genes [58]. AGO2 expression was induced by TCV infection [20]. In contrast, TuMV did not affect AGO accumulation [15]. Therefore, the availability and coordination among antiviral AGOs against different viruses could possibly be virus-dependent.

Furthermore, the capacity to physically interact with AGO protein was considered to be critical for the VSR functions of TCV CPs (P38) [59]. However, the VSR function of P37, coat protein of *Pelargonium line pattern virus,* was shown to be dependent on its sRNA binding capacity instead of its physical interactions with AGO protein [60]. Here, we showed that multiple AGOs participate in anti-TCV silencing. More molecular evidence is needed to understand how P38 interacts with these AGOs to suppress antiviral silencing machinery.

Regardless of precise antiviral mechanisms in play against TCV, the image-based phenotyping system developed here was shown to be useful in delineating both major and minor contributions of AGOs during virus infection over time. This should encourage development and refinement of additional tools and readouts to measure more effects and responses, even before visible symptoms of disease are detectable. Closing the gap between knowledge at the genetic and molecular levels with phenotypic effects during pathogen infection and non-pathogen colonization should yield considerable new insights into host-microbe interactions.

## Conclusion

Precise phenotyping methods to measure biotic and abiotic effects on plant health are important to develop. A high-throughput, image-based disease trait phenotyping pipeline to quantify virus-induced symptoms in *Arabidopsis* was developed. Combined with infectivity experiments using both parental and VSR-defective TCV variants, AGO2 and AGO7 were identified as the most prominent antiviral AGO proteins against TCV, while AGO3 was found to have a minor effect on antiviral silencing. These and previous data support the idea that multiple AGOs are recruited and programmed during antiviral defense in unique ways against different virus species.

## Materials and Methods

### Plant materials

All *Arabidopsis* plants used in this study (including all mutant lines) were in the *Columbia-0 (Col-0)* background and were grown in growth chambers under long day (16-hour light/8-hour dark) conditions, at 22°C and 75% relative humidity. The ten *ago* mutants used in this study were described previously: *ago1-27, ago2-1, ago3-2, ago4-2, ago5-2, ago6-3, zip-1, ago8-1, ago9-5* and *ago10-5* [15]. Double and triple *ago* mutants were generated by crossing between single *ago* mutants. The *dcl2-1 dcl3-1 dcl4-2* triple mutant was previously described [61].

*Nicotiana benthamiana* plants were grown in growth chambers under long day (16-hour light/8-hour dark) conditions, at 22°C and 75% relative humidity.

### DNA plasmids

The pSW-TCV plasmid is equivalent to pPZP212-TCV described before [26] (pSW is derived from pPZP212 plasmid by deleting most of RE sites with its MCS). The pSW-TCV CPB construct incorporates the mutation (R130T) in pSW-TCV CP region [26]. TCV CP fragment was amplified from pSW-TCV and pSW-TCV CPB using one set of primers: P38_L (ATGGAAAATGATCCTAGAGTCCG) and P38_R (CTAAATTCTGAGTGCTTGCCATTT). P38 PCR segments were sent to GENEWIZ to confirm the CPB mutation by using sequencing primer P38_L (ATGGAAAATGATCCTAGAGTCCG).

### Virus infection assays

Plasmids carrying TCV DNA clones were transformed into *Agrobacterium tumefaciens* (GV3101) and then infiltration was done on *N. benthamiana* leaves for viral inoculum preparation. The infected *N. benthamiana* leaves were grinded in 200 mM NaOAc solution (pH5.2) at 4°C. The grinded mixtures were centrifuged at 4°C to collect the supernatant. Next, 40% PEG-8000 (Polyethylene glycol; MW:8000) in 1M NaCl solution was added to the supernatant and incubated on ice overnight. Virion pellet were collected by centrifuging and resuspended in 10mM NaOAc (pH5.5) solution for inoculum stock. The inoculum stocks were aliquoted into equal volumes and stored at − 80°C. The same set of inoculums (1:10 dilution with 10mM NaOAc pH5.5) were used to inoculate all genotypes in this study. The *A. thaliana* plants to be inoculated were approximately two weeks old. The *ago1-27* mutant plant was planted one week earlier than other genotypes. The four largest rosette leaves were inoculated by gently rubbing carborundum dusted on the leaf surface. Plants under different treatments were placed on separate benches to avoid possible cross contaminations in the growth chamber.

### Protein blot assays

To measure local CP accumulation, four inoculated rosette leaves per plant were collected at 7 days post inoculation (dpi) and pooled into a single sample. For systemic CP accumulation, the four largest non-inoculated cauline leaves or five to six inflorescence clusters per plant were collected at 14 dpi and pooled into a single sample. Four biological replicates were randomly collected from each genotype-treatment group. Total protein was extracted and normalized to 0.5 μg/μl. Protein samples (6μg each) were separated on NuPAGE 4–12% Bis-Tris Protein gels and subsequently transferred to nitrocellulose blotting members (0.45μm, GE Healthcare Life Science) for protein detection with corresponding antibodies. TCV CP was detected using anti-TCV-P38 serum (F. Qu, personal communication) at a dilution of 1:20,000 and the large subunit of rubisco protein was detected using anti-rubisco (plant) antibody in chicken (Sigma-Aldrich) at 1:10,000. The blots were incubated with goat anti-rabbit lgG-HRP conjugate (GE Healthcare) secondary serum at a dilution of 1:10,000 to detect TCV CP and with rabbit anti-chicken lgY (H+L) HRP conjugate secondary antibody (Thermo Fisher) at a dilution of 1: 10,000 to detect rubisco protein. TCV CP and rubisco protein were detected on the same blot. Quantification of western blot images was done by normalizing all other bands relative to *dcl2-1 dcl3-1 dcl4-2* (P38 band/ loading control band) using *ImageJ* software [60].

### Manual plant height measurement

Measurement of plant height was done manually using a cubic ruler to measure from the rosette plane to the top of the main plant stem [62]. Data were collected every other day from the day of virus inoculation to 21 dpi.

### RGB Image acquisition

The soil in the growth pots was covered by a blue mesh (commercially available from Amazon.com), leaving a hole in the center for the plant to grow out (Fig 2B). Meshed plants were bottom-watered to avoid water splash on plant tissues. Images of plant rosettes were manually acquired using a Canon Digital Rebel XT DSLR camera with EF-S 18–55mm f2.5-5.6 lens (0.60 s exposure, F/14, ISO100) on a Kaiser RS1 Copy Stand. The flash function was kept off and fixed ambient light was used in a closed room to minimize illumination discrepancies between images. An X-rite ColorChecker Digital SG was placed next to the plant as a color reference and correction. Images were stored in the native RAW format and also high-resolution JPG format. Plants were removed from the growth chamber (10 am −11 am) and any bolts above the rosette plane were removed before imaging. Stationary plant rosettes were imaged every other day from one day before inoculation to 17 dpi.

### Image Analysis

Color images of individual mock- or virus-infected *Arabidopsis* plants were processed using the Plant Computer Vision (PlantCV) package to quantify the progression of disease symptoms [32, 33]. PlantCV v2.1 (commit: d553a2c1c6bd29e734d19898e3e9ac4fcff40aa9) was used. Image analysis was done in parallel using HTCondor v8.6.8 [63].

The PlantCV naive Bayes machine learning method [31, 32] was used to segment image pixels into four classes: 1) green plant pixels (“healthy”); 2) chlorotic/necrotic plant pixels (“unhealthy”); 3) background pixels from the blue mesh material used to cover the soil, and 4) all other background pixels. The PlantCV naive Bayes classifier was trained using pixel red, green, blue (RGB) color values from 1980 and 3779 pixels manually sampled from the background and foreground classes, respectively, from multiple images using the ImageJ pixel picking tool [64] as described in [32]. The training data was used to calculate probability density functions (PDFs) using kernel density estimation (KDE) for each class in the hue, saturation, and value (HSV) color space [32]. During image analysis, the PDFs were used to parameterize the naive Bayes classifier to segment images into the four output classes (one binary image per class).

After segmentation, the blue mesh was used to automatically identify the position of the pot within each image. Morphological erosion was used to reduce noise in the classified blue mesh pixels. A universal region of interest was used to keep blue mesh connected components in the center area of each image since the pot was consistently centered in each image. The remaining blue mesh connected components were flattened into a single object and were used to create a pot binary mask using the bounding rectangle area of the blue mesh. Padding was added to the minimum bounding rectangle area using morphological dilation. The resulting pot binary mask was used to mask the “healthy” and “unhealthy” binary masks to remove misclassified background pixels. The “healthy” and “unhealthy” binary images were further filtered to remove background pixels misclassified as foreground pixels using a size-based filter that removed small connected components (300 and 1000 pixels for “healthy” and “unhealthy” binary images, respectively). A universal region of interest was used to keep connected components in the center area of each image since the Arabidopsis plants were consistently centered in each image. The resulting cleaned binary image for the “healthy” and “unhealthy” classes were used to measure the area of green and chlorotic/necrotic pixels in each image.

Additionally, the union of the “healthy” and “unhealthy” plant pixels was calculated to create a combined plant mask. The input RGB images were converted to HSV color space and the frequency distribution of hue color values for each plant at each timepoint was extracted from the pixels that overlapped the combined plant mask. The hue color histograms were plotted in R v3.4.4 using the libraries ggplot2, ggridges, reshape2, and dplyr [65–69]. The PlantCV and R analysis code and result files are available at GitHub (https://github.com/carringtonlab/tcv-image-analysis).

### Statistics

All statistical analysis was performed in R v3.4.4.

## Acknowledgments

We are grateful for Dr. Feng Qu (Department of Plant Pathology, The Ohio State University, USA) for providing pSW-TCV plasmids and anti-P38 antibody. We thank Kerrigan B. Gilbert, Dr. Dan Lin, Dr. Steen Hoyer and Dr. Kira Veley for support and critique of the study. We also thank Robyn Allscheid for constructive editorial advices on this manuscript.

The authors have declared no conflicts of interests.

This study was supported by funding: National Institutes of Health (www.nih.gov) grant (AI043288) to JCC. National Science Foundation (www.nsf.gov) grant (1330562) to JCC.

## Author Contributions

X.Z. and J.C.C conceived and designed research. X.Z. and A.A. performed experiments. X.Z., N.F. and J.B. analyzed and plotted data. J.C.C. and X.Z. wrote the manuscript with contribution of all authors.

## Supplemental Data

**S1 Fig. CPB mutation in TCV coat protein does not affect its function in virion assembly.**

(A) TCV virions extracted from *Col-0* or *dcl2 dcl3 dcl4* plants inoculated with TCV or TCV CPB were stained with Ethidium Bromide (left panel) for viral genomic RNAs or Coomassie Blue (right panel) for viral coat proteins. Virions were extracted from pooled systemic non-inoculated leaves at 14 dpi. Two biological replicates were shown in the same gel. (B) TCV virions from TCV or TCV CPB inoculum with different dilutions were stained with Ethidium Bromide (left panel) for viral genomic RNAs or Coomassie Blue (right panel) for viral coat proteins. Virions extracted from *Col-0* inoculated rosette leaves (RL) and non-inoculated cauline leaves (CL) infected with TCV were loaded and stained as positive controls.

**S2 Fig. Temporal visualization of rosette leaves and the corresponding pseudo-color images in *ago1–27* mutant.**

Plants were inoculated with mock solution, TCV or TCV CPB inoculum separately. Photos were taken from 1 day before inoculation to 17 days post inoculation (dpi). In the pseudo-color images, green color referred to healthy plant pixels, while red color referred to chlorotic/necrotic plant pixels.

**S3 Fig. Temporal visualization of rosette leaves and the corresponding pseudo-color images in *ago2–1* mutant.**

Plants were inoculated with mock solution, TCV or TCV CPB inoculum separately. Photos were taken from 1 day before inoculation to 17 days post inoculation (dpi). In the pseudo-color images, green color referred to healthy plant pixels, while red color referred to chlorotic/necrotic plant pixels.

**S4 Fig. Temporal visualization of rosette leaves and the corresponding pseudo-color images in *ago3–2* mutant.**

Plants were inoculated with mock solution, TCV or TCV CPB inoculum separately. Photos were taken from 1 day before inoculation to 17 days post inoculation (dpi). In the pseudo-color images, green color referred to healthy plant pixels, while red color referred to chlorotic/necrotic plant pixels.

**S5 Fig. Temporal visualization of rosette leaves and the corresponding pseudo-color images in *ago4–2* mutant.**

Plants were inoculated with mock solution, TCV or TCV CPB inoculum separately. Photos were taken from 1 day before inoculation to 17 days post inoculation (dpi). In the pseudo-color images, green color referred to healthy plant pixels, while red color referred to chlorotic/necrotic plant pixels.

**S6 Fig. Temporal visualization of rosette leaves and the corresponding pseudo-color images in *ago5–2* mutant.**

Plants were inoculated with mock solution, TCV or TCV CPB inoculum separately. Photos were taken from 1 day before inoculation to 17 days post inoculation (dpi). In the pseudo-color images, green color referred to healthy plant pixels, while red color referred to chlorotic/necrotic plant pixels.

**S7 Fig. Temporal visualization of rosette leaves and the corresponding pseudo-color images in *ago6-3* mutant.**

Plants were inoculated with mock solution, TCV or TCV CPB inoculum separately. Photos were taken from 1 day before inoculation to 17 days post inoculation (dpi). In the pseudo-color images, green color referred to healthy plant pixels, while red color referred to chlorotic/necrotic plant pixels.

**S8 Fig. Temporal visualization of rosette leaves and the corresponding pseudo-color images in *zip-1* mutant.**

Plants were inoculated with mock solution, TCV or TCV CPB inoculum separately. Photos were taken from 1 day before inoculation to 17 days post inoculation (dpi). In the pseudo-color images, green color referred to healthy plant pixels, while red color referred to chlorotic/necrotic plant pixels.

**S9 Fig. Temporal visualization of rosette leaves and the corresponding pseudo-color images in *ago8-5* mutant.**

Plants were inoculated with mock solution, TCV or TCV CPB inoculum separately. Photos were taken from 1 day before inoculation to 17 days post inoculation (dpi). In the pseudo-color images, green color referred to healthy plant pixels, while red color referred to chlorotic/necrotic plant pixels.

**S10 Fig. Temporal visualization of rosette leaves and the corresponding pseudo-color images in *ago9-5* mutant.**

Plants were inoculated with mock solution, TCV or TCV CPB inoculum separately. Photos were taken from 1 day before inoculation to 17 days post inoculation (dpi). In the pseudo-color images, green color referred to healthy plant pixels, while red color referred to chlorotic/necrotic plant pixels.

**S11 Fig. Temporal visualization of rosette leaves and the corresponding pseudo-color images in *ago10-5* mutant.**

Plants were inoculated with mock solution, TCV or TCV CPB inoculum separately. Photos were taken from 1 day before inoculation to 17 days post inoculation (dpi). In the pseudo-color images, green color referred to healthy plant pixels, while red color referred to chlorotic/necrotic plant pixels.

**S12 Fig. Temporal color change in rosette leaves in ten single *ago* mutant plants infected by TCV.**

Each histogram illustrated the distribution of pixels at each hue value (degree) in a single genotype and treatment combination at different time point. The temporal change of pixel distribution could be observed from 1 day before inoculation to 17 dpi. Pixels were collected and summed from four individual images in the same genotype and treatment combination: mock (left panel), TCV (middle panel), or TCV CPB (right panel) inoculum. The color chart illustrates the relationship between the hue value (degree) with the corresponding RGB coordinates.

**S1 Table. Kolmogorov-Smirnov statistical results for rosette size curve in single *ago* mutants.**

**S2 Table. Kolmogorov-Smirnov statistical results for the percentage of unhealthy tissue curve in single *ago* mutants.**

**S3 Table. Kolmogorov-Smirnov statistical results for average hue curve in single *ago* mutants.**

**S4 Table. Kolmogorov-Smirnov statistical results for plant height curve in single *ago* mutants.**

## References

1. Agius, C., et al., RNA silencing and antiviral defense in plants. Methods Mol Biol, 2012. 894: p. 17–38.

2. Bologna, N.G. and O. Voinnet, The diversity, biogenesis, and activities of endogenous silencing small RNAs in Arabidopsis. Annu Rev Plant Biol, 2014. 65: p. 473–503.

3. Fang, X. and Y. Qi, RNAi in Plants: An Argonaute-Centered View. Plant Cell, 2016. 28(2): p. 272–85.

4. Incarbone, M. and P. Dunoyer, RNA silencing and its suppression: novel insights from in planta analyses. Trends Plant Sci, 2013. 18(7): p. 382–92.

5. Bortolamiol, D., et al., The Polerovirus F box protein P0 targets ARGONAUTE1 to suppress RNA silencing. Curr Biol, 2007. 17(18): p. 1615–21.

6. Pazhouhandeh, M., et al., F-box-like domain in the polerovirus protein P0 is required for silencing suppressor function. Proc Natl Acad Sci U S A, 2006. 103(6): p. 1994–9.

7. Chiu, M.H., et al., The silencing suppressor P25 of Potato virus X interacts with Argonaute1 and mediates its degradation through the proteasome pathway. Mol Plant Pathol, 2010. 11(5): p. 641–9.

8. Baumberger, N., et al., The Polerovirus silencing suppressor P0 targets ARGONAUTE proteins for degradation. Curr Biol, 2007. 17(18): p. 1609–14.

9. Zhang, X., et al., Cucumber mosaic virus-encoded 2b suppressor inhibits Arabidopsis Argonaute1 cleavage activity to counter plant defense. Genes Dev, 2006. 20(23): p. 3255–68.

10. Fang, Y.Y., et al., CMV2b-AGO Interaction Is Required for the Suppression of RDR-Dependent Antiviral Silencing in Arabidopsis. Front Microbiol, 2016. 7: p. 1329.

11. Giner, A., et al., Viral protein inhibits RISC activity by argonaute binding through conserved WG/GW motifs. PLoS Pathog, 2010. 6(7): p. e1000996.

12. Kenesi, E., et al., A viral suppressor of RNA silencing inhibits ARGONAUTE 1 function by precluding target RNA binding to pre-assembled RISC. Nucleic Acids Res, 2017. 45(13): p. 7736–7750.

13. Varallyay, E., et al., Plant virus-mediated induction of miR168 is associated with repression of ARGONAUTE1 accumulation. EMBO J, 2010. 29(20): p. 3507–19.

14. Qu, F., X. Ye, and T.J. Morris, Arabidopsis DRB4, AGO1, AGO7, and RDR6 participate in a DCL4-initiated antiviral RNA silencing pathway negatively regulated by DCL1. Proc Natl Acad Sci U S A, 2008. 105(38): p. 14732–7.

15. Garcia-Ruiz, H., et al., Roles and programming of Arabidopsis ARGONAUTE proteins during Turnip mosaic virus infection. PLoS Pathog, 2015. 11(3): p. e1004755.

16. Carbonell, A. and J.C. Carrington, Antiviral roles of plant ARGONAUTES. Curr Opin Plant Biol, 2015. 27: p. 111–7.

17. Wang, X.B., et al., The 21-nucleotide, but not 22-nucleotide, viral secondary small interfering RNAs direct potent antiviral defense by two cooperative argonautes in Arabidopsis thaliana. Plant Cell, 2011. 23(4): p. 1625–38.

18. Dzianott, A., J. Sztuba-Solinska, and J.J. Bujarski, Mutations in the antiviral RNAi defense pathway modify Brome mosaic virus RNA recombinant profiles. Mol Plant Microbe Interact, 2012. 25(1): p. 97–106.

19. Kontra, L., et al., Distinct Effects of p19 RNA Silencing Suppressor on Small RNA Mediated Pathways in Plants. PLoS Pathog, 2016. 12(10): p. e1005935.

20. Harvey, J.J., et al., An antiviral defense role of AGO2 in plants. PLoS One, 2011. 6(1): p. e14639.

21. Jaubert, M., et al., ARGONAUTE2 mediates RNA-silencing antiviral defenses against Potato virus X in Arabidopsis. Plant Physiol, 2011. 156(3): p. 1556–64.

22. Scholthof, H.B., et al., Identification of an ARGONAUTE for antiviral RNA silencing in Nicotiana benthamiana. Plant Physiol, 2011. 156(3): p. 1548–55.

23. Raja, P., et al., Arabidopsis double-stranded RNA binding protein DRB3 participates in methylation-mediated defense against geminiviruses. J Virol, 2014. 88(5): p. 2611–22.

24. Raja, P., et al., Viral genome methylation as an epigenetic defense against geminiviruses. J Virol, 2008. 82(18): p. 8997–9007.

25. Cohen, Y., A. Gisel, and P.C. Zambryski, Cell-to-cell and systemic movement of recombinant green fluorescent protein-tagged turnip crinkle viruses. Virology, 2000. 273(2): p. 258–66.

26. Cao, M., et al., The capsid protein of Turnip crinkle virus overcomes two separate defense barriers to facilitate systemic movement of the virus in Arabidopsis. J Virol, 2010. 84(15): p. 7793–802.

27. Donze, T., et al., Turnip crinkle virus coat protein inhibits the basal immune response to virus invasion in Arabidopsis by binding to the NAC transcription factor TIP. Virology, 2014. 449: p. 207–14.

28. Chen, Y.J., et al., The capsid protein p38 of turnip crinkle virus is associated with the suppression of cucumber mosaic virus in Arabidopsis thaliana co-infected with cucumber mosaic virus and turnip crinkle virus. Virology, 2014. 462–463: p. 71–80.

29. Thomas, C.L., et al., Turnip crinkle virus coat protein mediates suppression of RNA silencing in Nicotiana benthamiana. Virology, 2003. 306(1): p. 33–41.

30. Bock, C.H., et al., Plant Disease Severity Estimated Visually, by Digital Photography and Image Analysis, and by Hyperspectral Imaging. Critical Reviews in Plant Sciences, 2010. 29(2): p. 59–107.

31. Abbasi, A. and N. Fahlgren, Naïve Bayes pixel-level plant segmentation, in 2016 IEEE Western New York Image and Signal Processing Workshop (WNYISPW). 2016. p. 1–4.

32. Gehan, M.A., et al., PlantCV v2: Image analysis software for high-throughput plant phenotyping. PeerJ, 2017. 5: p. e4088.

33. Fahlgren, N., et al., A Versatile Phenotyping System and Analytics Platform Reveals Diverse Temporal Responses to Water Availability in Setaria. Mol Plant, 2015. 8(10): p. 1520–35.

34. Meng, C., et al., Host-induced avirulence of hibiscus chlorotic ringspot virus mutants correlates with reduced gene-silencing suppression activity. J Gen Virol, 2006. 87(Pt 2): p. 451–9.

35. Martinez-Turino, S. and C. Hernandez, Inhibition of RNA silencing by the coat protein of Pelargonium flower break virus: distinctions from closely related suppressors. J Gen Virol, 2009. 90(Pt 2): p. 519–25.

36. Sass, L., P. Majer, and E. Hideg, Leaf hue measurements: a high-throughput screening of chlorophyll content. Methods Mol Biol, 2012. 918: p. 61–9.

37. Liang, Y., et al., A nondestructive method to estimate the chlorophyll content of Arabidopsis seedlings. Plant Methods, 2017. 13: p. 26.

38. Veley, K.M., et al., High-throughput profiling and analysis of plant responses over time to abiotic stress. Plant Direct, 2017(1): p. 4.

39. Gonzalez RC, W.R., Eddins SL, Digital image processing using MATLAB. 2009, New York: Gatesmark Publishing.

40. Boyes, D.C., et al., Growth stage-based phenotypic analysis of Arabidopsis: a model for high throughput functional genomics in plants. Plant Cell, 2001. 13(7): p. 1499–510.

41. Nguyen, H.P., et al., Methods to Study PAMP-Triggered Immunity Using Tomato and Nicotiana benthamiana. Molecular Plant-Microbe Interactions, 2010. 23(8): p. 991–999.

42. Arend, D., et al., Quantitative monitoring of Arabidopsis thaliana growth and development using high-throughput plant phenotyping. Scientific Data, 2016. 3.

43. Lloyd, S.R., et al., Methods to Study PAMP-Triggered Immunity in Brassica Species. Molecular Plant-Microbe Interactions, 2014. 27(3): p. 286–295.

44. Laflamme, B., et al., Image-Based Quantification of Plant Immunity and Disease. Molecular Plant-Microbe Interactions, 2016. 29(12): p. 919–924.

45. Pethybridge, S.J. and S.C. Nelson, Leaf Doctor: A New Portable Application for Quantifying Plant Disease Severity. Plant Disease, 2015. 99(10): p. 1310–1316.

46. Mutka, A.M. and R.S. Bart, Image-based phenotyping of plant disease symptoms. Frontiers in Plant Science, 2015. 5.

47. Singh, A., et al., Machine Learning for High-Throughput Stress Phenotyping in Plants. Trends in Plant Science, 2016. 21(2): p. 110–124.

48. Lee, U., et al., An automated, high-throughput plant phenotyping system using machine learning based plant segmentation and image analysis. Plos One, 2018. 13(4).

49. Zhang, Z.H., et al., Arabidopsis AGO3 predominantly recruits 24-nt small RNAs to regulate epigenetic silencing. Nature Plants, 2016. 2(5).

50. Schuck, J., et al., AGO/RISC-mediated antiviral RNA silencing in a plant in vitro system. Nucleic Acids Research, 2013. 41(9): p. 5090–5103.

51. Minoia, S., et al., Specific Argonautes Selectively Bind Small RNAs Derived from Potato Spindle Tuber Viroid and Attenuate Viroid Accumulation In Vivo. Journal of Virology, 2014. 88(20): p. 11933–11945.

52. Montgomery, T.A., et al., Specificity of ARGONAUTE7-miR390 interaction and dual functionality in TAS3 trans-acting siRNA formation. Cell, 2008. 133(1): p. 128–141.

53. Axtell, M.J., et al., A two-hit trigger for siRNA biogenesis in plants. Cell, 2006. 127(3): p. 565–577.

54. Axtell, M.J., J.A. Snyder, and D.P. Bartell, Common functions for diverse small RNAs of land plants. Plant Cell, 2007. 19(6): p. 1750–1769.

55. Jouannet, V., et al., Cytoplasmic Arabidopsis AGO7 accumulates in membrane-associated siRNA bodies and is required for ta-siRNA biogenesis. Embo Journal, 2012. 31(7): p. 1704–1713.

56. Carbonell, A., et al., Functional Analysis of Three Arabidopsis ARGONAUTES Using Slicer-Defective Mutants. Plant Cell, 2012. 24(9): p. 3613–3629.

57. Burgyan, J. and Z. Havelda, Viral suppressors of RNA silencing. Trends in Plant Science, 2011. 16(5): p. 265–272.

58. Wu, C., et al., Analyses of RNA-Seq and sRNA-Seq data reveal a complex network of anti-viral defense in TCV-infected Arabidopsis thaliana. Scientific Reports, 2016. 6.

59. Azevedo, J., et al., Argonaute quenching and global changes in Dicer homeostasis caused by a pathogen-encoded GW repeat protein. Genes & Development, 2010. 24(9): p. 904–915.

60. Perez-Canamas, M. and C. Hernandez, Key Importance of Small RNA Binding for the Activity of a Glycine-Tryptophan (GW) Motif-containing Viral Suppressor of RNA Silencing. Journal of Biological Chemistry, 2015. 290(5): p. 3106–3120.

61. Deleris, A., et al., Hierarchical action and inhibition of plant Dicer-like proteins in antiviral defense. Science, 2006. 313(5783): p. 68–71.

62. Boyes, D.C., et al., Growth stage-based phenotypic analysis of arabidopsis: A model for high throughput functional genomics in plants. Plant Cell, 2001. 13(7): p. 1499–1510.

63. Thain, D., T. Tannenbaum, and M. Livny, Distributed computing in practice: the Condor experience. Concurrency and Computation-Practice & Experience, 2005. 17(2–4): p. 323–356.

64. Schneider, C.A., W.S. Rasband, and K.W. Eliceiri, NIH Image to ImageJ: 25 years of image analysis. Nature Methods, 2012. 9(7): p. 671–675.

65. Wickham, H., Reshaping data with the reshape package. Journal of Statistical Software, 2007. 21(12): p. 1–20.

66. Wickham H F.R., Henry L, Müller K. dplyr: A Grammar of Data Manipulation. 2017; Available from: https://dplyr.tidyverse.org/.

67. Wickham, H., ggplot2: Elegant Graphics for Data Analysis. 2009: Springer-Verlag New York.

68. Wilke, C.O. ggridges: Ridgeline Plots in ‘ggplots’. 2017; Available from: https://cran.r-project.org/web/packages/ggridges/vignettes/introduction.html.

69. Team, R.C. R: A Language and Environment for Statistical Computing. 2018; Available from: https://www.r-project.org/.

